# SNAP: Streamlined Nextflow Analysis Pipeline for Immunoprecipitation-Based Epigenomic Profiling of Circulating Chromatin

**DOI:** 10.64898/2025.12.30.694452

**Authors:** Ze Zhang, Paulo Da Silva Cordeiro, Surya B. Chhetri, Brad Fortunato, Zhenjie Jin, Razane El Hajj Chehade, Karl Semaan, Gunsagar Gulati, Garyoung Gary Lee, Christopher Hemauer, Weiwei Bian, Shahabeddin Sotudian, Ziwei Zhang, David Osei-Hwedieh, Tanya E. Heim, Corrie Painter, Rashad Nawfal, Marc Eid, Damien Vasseur, John Canniff, Hunter Savignano, Noa Phillips, Ji-Heui Seo, Kurt R. Weiss, Matthew L. Freedman, Sylvan C. Baca

## Abstract

Epigenomic profiling of circulating chromatin is a powerful and minimally invasive approach for detecting and monitoring disease, but there are no bioinformatics pipelines tailored to the unique characteristics of cell-free chromatin. We present SNAP (Streamlined Nextflow Analysis Pipeline), a reproducible, scalable, and modular workflow specifically designed for immunoprecipitation-based methods for profiling cell-free chromatin. SNAP incorporates quality control metrics optimized for circulating chromatin, including enrichment score and fragment count thresholds, as well as direct estimation of circulating tumor DNA (ctDNA) content from fragment length distributions. It also includes SNP fingerprinting to enable sample identity verification. When applied to cfChIP-seq and cfMeDIP-seq data across multiple cancer types, SNAP’s quality filters significantly improved classification performance while maintaining high data retention. Independent validation using plasma from patients with osteosarcoma confirmed the detection of tumor-associated epigenomic signatures that correlated with ctDNA levels and reflected disease biology. SNAP’s modular architecture enables straightforward extension to additional cell-free immunoprecipitation-based assays, providing a robust framework to support studies of circulating chromatin broadly. SNAP is compatible with cloud and high-performance computing environments and is publicly available at https://github.com/prc992/SNAP/.

**Graphic Abstract:** 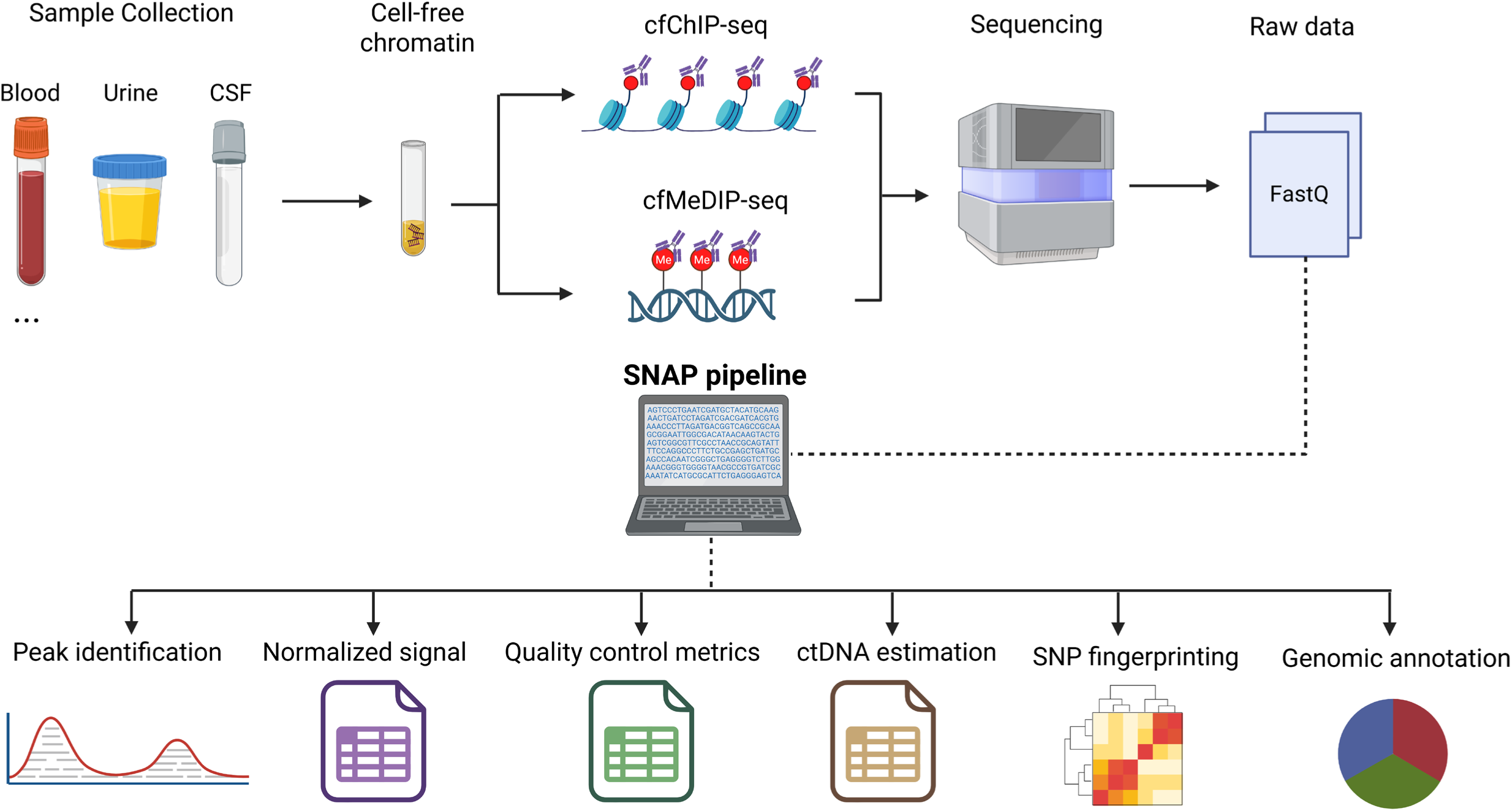

## Introduction

Liquid biopsy is increasingly used to inform clinical decision-making in precision medicine, offering a scalable and noninvasive platform for disease detection, longitudinal monitoring, and treatment stratification^1–3^. Profiling circulating chromatin released into biofluids such as blood and urine provides insights into tissue-of-origin, cell states, and gene regulation^4–8^. Chromatin features associated with promoters, enhancers, and gene bodies serve as robust proxies for transcriptional activity, enabling inference of gene regulatory programs that reflect underlying disease biology and its dynamics^9–11^.

Immunoprecipitation (IP)-based approaches such as cell-free chromatin immunoprecipitation sequencing (cfChIP-seq) and cell-free methylated DNA immunoprecipitation sequencing (cfMeDIP-seq) enable genome-wide profiling of histone modifications and DNA methylation in circulating chromatin. For example, cfChIP-seq can measure histone modifications associated with active gene promoters (H3K4me3), promoters and enhancers (H3K27ac), and transcribed gene bodies (H3K36me3). cfMeDIP-seq can detect repressive DNA methylation at CpG islands as well as lineage-associated patterns of DNA methylation. An advantage of these IP-based approaches is that they enrich for epigenomic features of interest, which can improve signal-to-noise ratios thereby decreasing the amount of DNA sequencing required for other approaches.

cfChIP-seq and cfMeDIP-seq have been applied across a range of conditions, including various cancer types^7–10,12–17^, organ transplants^18–20^, autoimmune diseases^21,22^, reproductive health^23^, infectious diseases^24,25^, and prenatal screening^26^, highlighting their broad applicability and potential for precision medicine. Biofluids beyond blood plasma are increasingly being profiled, including cerebrospinal fluid (CSF)^15^, ovarian follicular fluid^23^, and urine^27^. Circulating chromatin profiling has become a focus of intense investigation, with emerging technologies fueling the generation of thousands of datasets over the last few years. This rapid expansion underscores the need for standardized, scalable tools to analyze and interpret these data.

Despite this growing interest, dedicated pipelines for analyzing cfChIP-seq and cfMeDIP-seq data remain lacking. Most existing studies have relied on project-specific, *ad hoc* scripts for read processing, quality assessment, and signal quantification, limiting reproducibility and cross-study comparisons^9,10^. Moreover, the circulating chromatin poses analytical challenges, such as low-input material, high fragmentation, and the need for quantitative signal interpretation. Conventional ChIP-seq pipelines, which are primarily focused on identifying the presence of “peaks” where signal is elevated above background, are poorly suited to cfChIP-seq and cfMeDIP-seq data^28–30^. In addition, ChIP-seq quality control metrics such as peak number are not optimized for cell-free contexts, and normalization strategies tailored to the unique properties of these assays are critically needed^31–33^.

To address these challenges, we developed SNAP, a modular and reproducible workflow specifically designed for IP-based profiling of circulating chromatin. Built on the Nextflow framework and Docker containerization, SNAP ensures reproducibility, portability across computing environments, and scalability through efficient parallelization. The pipeline incorporates assay-specific quality control metrics, enrichment scoring to assess immunoprecipitation specificity, signal normalization strategies using housekeeping regions, and estimation of tumor DNA fraction based on fragment lengths^34^. It also integrates SNP fingerprinting for sample identity verification^35^, a critical step when profiling multiple epigenetic features from one sample or multiple longitudinal samples from the same subject to guard against sample swaps. As the generation of cfChIP-seq and cfMeDIP-seq data accelerates, SNAP offers a standardized, scalable, and biologically meaningful framework for analyzing circulating chromatin across diverse contexts in human health and disease.

## Methods

### Pipeline Architecture and Development

SNAP is a comprehensive computational framework designed specifically for analyzing epigenetic modifications in circulating chromatin (**Figure 1A**). Built using Nextflow^36^, a domain-specific language, the pipeline ensures reproducibility, scalability, and portability across diverse computing environments. The architecture follows a modular design pattern where each processing step is implemented within a distinct module, facilitating flexible execution and seamless maintenance (**Figure 1B**). The pipeline employs a sub-workflow architecture that organizes related processes into modular units, allowing users to start from different entry points (e.g., FASTQ or BAM inputs) and to stop execution at any desired stage. When execution is not completed, cumulative intermediate reports are still generated, summarizing all steps performed up to that point. This design can accommodate varying research requirements and computational constraints while maintaining a structured analytical workflow.

**Figure 1.**
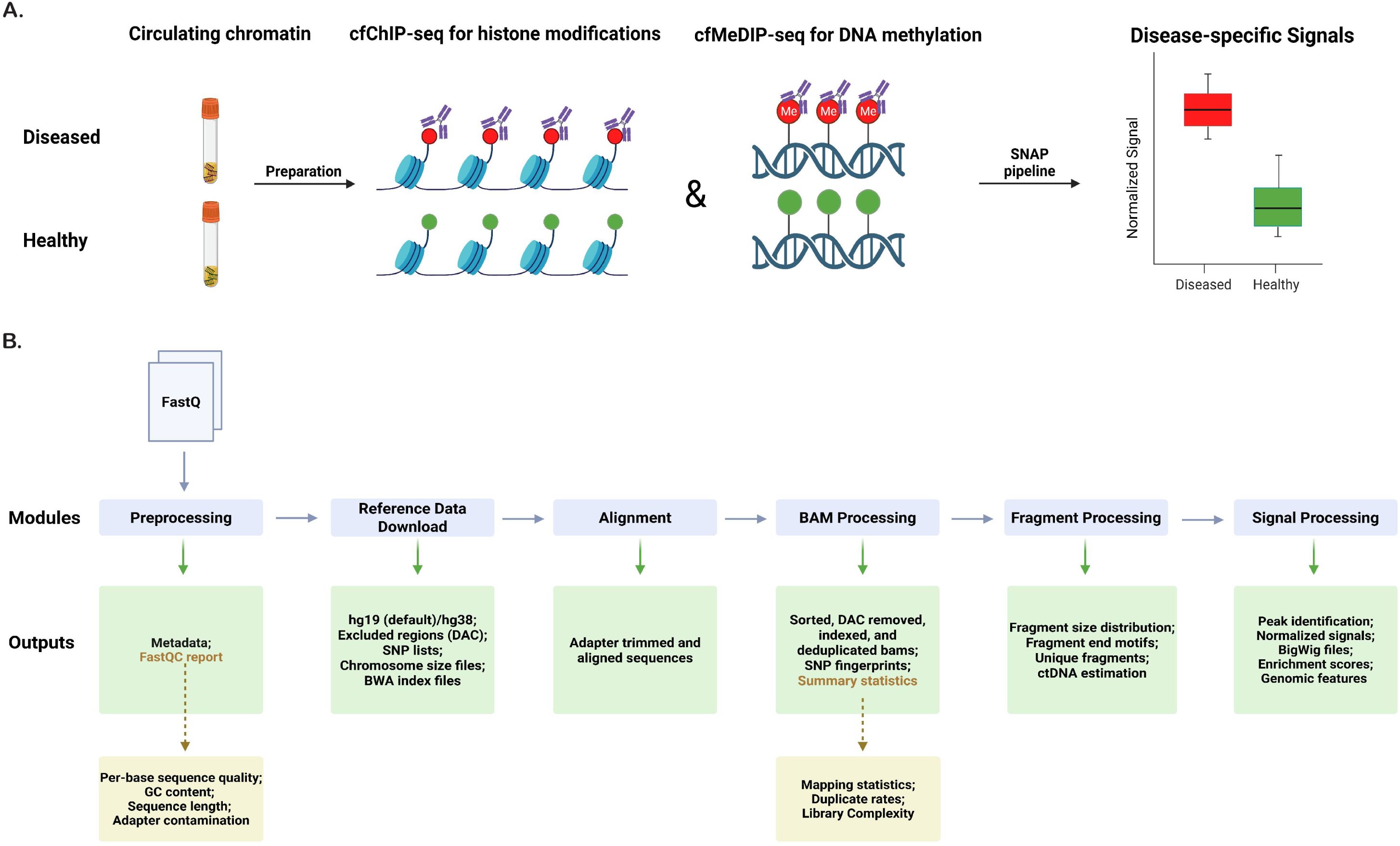
Overview of the SNAP pipeline. **(A).** Biological rationale for applying SNAP to plasma cfDNA epigenomic profiling. **(B).** Schematic of the SNAP computational workflow.

### Preprocessing

This module executes *FastQC* (v0.11.9) to generate quality control reports for raw sequencing data^37^. *FastQC* evaluates key quality metrics such as per-base sequence quality, GC content distribution, sequence length, and adapter contamination, which can identify sequencing quality issues that may affect the accuracy of downstream analyses. In addition, this module provides a convenient option for users who prefer to initiate the pipeline by simply specifying a directory containing all FASTQ files grouped by sample, instead of manually creating a sample sheet listing each sample and file path. In this mode, the *createSamplesheet* subroutine automatically scans the directory provided via the *--sample_dir_fasta* parameter and generates a spreadsheet (CSV format) with the following structure: *sampleId, enrichment_mark, read1, read2, control*.

### Reference Genome Management

The “reference download” sub-workflow manages multiple genome builds, with hg19 serving as the default reference. This module implements an automated system for downloading and preparing reference genomes while ensuring consistency across analysis runs. The process retrieves chromosome sizes, informative SNP lists required for sample fingerprinting^35^, and genomic exclusion regions defined by the ENCODE consortium^38^. Although time-intensive during initial execution, particularly when downloading references and building indices, users can circumvent this step by manually placing the required files into the */SNAP/ref_files/genome* directory. *BWA* (v0.7.18) performs genome indexing, generating the necessary files for the *BWA-MEM* algorithm, including BWA index files and the reference genome in FASTA format^39^. Additionally, the pipeline creates chromosome size files using UCSC tools, essential for bigWig generation and genomic region calculations^40^. Integration of ENCODE (DAC) exclusion regions increases analysis specificity by masking genomic loci prone to sequencing or mapping artifacts, such as highly repetitive or low-mappability regions that often generate spurious ChIP-seq signal enrichment^38^.

### Alignment

The “alignment” sub-workflow is one of the most computationally intensive components of the pipeline. Adapter trimming is performed using *FASTP* (v1.0.1) by default^41^, which applies stringent base-quality and adapter-removal filters equivalent to those implemented in *Trim Galore* (v0.6.7)^42^. Users may alternatively select *Trim Galore* via the *--trim_method* parameter, which is available for compatibility. Following trimming, *BWA-MEM* performs sequence alignment with optimized parameters (-I 250,200,2000,10) tailored to cfDNA characteristics, such as variable fragment lengths and occasional structural variants^39^. Trimming quality thresholds and fragment-size modeling options can be customized via the *trimming_params* and *bwa_params* variables in the *nextflow.config* file. The pipeline supports both paired-end (PE) and single-end (SE) sequencing data, applying distinct quality filters through *SAMtools* (v1.15.1) optimized for each data type (PE: -f 3 -F 3844 -q 30; SE: -F 3844 -q 30)^43^. These stringent filters retain only high-quality, properly mapped reads, minimizing non-specific signal and improving the reliability of downstream analyses.

### BAM Processing

The “BAM processing” sub-workflow implements essential steps to enhance the quality and reliability of aligned sequencing data. Following alignment, the module performs sorting and filtering to retain only properly paired reads with sufficient sequencing quality, while excluding noise-prone genomic regions, followed by comprehensive indexing. Duplicate fragment marking is carried out using *Picard MarkDuplicates* (v2.27.4), which identifies and flags PCR duplicates to prevent artificial inflation of coverage and ensure accurate quantification of unique DNA fragments^44^. Library complexity estimation is performed with *Preseq* (v2.0.2), which projects the number of unique molecules and helps assess whether sequencing depth is sufficient for capturing most DNA molecules in a library^45^. As noted in the *Picard* documentation, low library complexity can arise from insufficient starting material or over-amplification^44^. In cfDNA-based assays like cfChIP-seq, these conditions are common due to the limited and fragmented nature of circulating DNA, which often results in higher duplication rates and reduced library diversity compared to conventional ChIP-seq experiments. A distinctive feature of this sub-workflow is the SNP fingerprinting analysis performed with *SMaSH*^35^, which constructs dendrograms for sample identification, verifying sample identity, and detecting potential contamination or sample mix-ups. Alignment statistics generation through *Samtools* produces comprehensive quality metrics^43^, including mapping statistics, duplicate rates, and library complexity estimates, crucial for assessing overall sequencing data quality.

### Fragment Analysis and Characterization

The “fragment processing” module performs detailed DNA fragment analysis, which can be used to derive biological insights from cfDNA. Fragment size distributions are visualized to confirm nucleosomal digestion patterns expected in cfDNA and to potentially flag contamination with genomic DNA that has distinct size distributions. The module extracts features that can be input into downstream fragmentomics analyses, including fragment end motifs, sizes, and GC content. Fragment locations are output into a bed file. Additionally, this module estimates ctDNA content from fragment sizes using Fragle^34^.

### Signal Processing and Peak Identification

The “signal processing” sub-workflow implements signal analysis and peak calling. BigWig and bedGraph file generation facilitates visualization with Integrative Genomics Viewer (IGV)^46^ or other visualization tools. This module produces IGV session files allowing rapid manual inspection of relevant regions defined in *ref_files/pileup_report/regions_of_interest.bed*. Peak calling employs *MACS2* (v2.2.7.1)^47^, specifically designed for ChIP-seq data analysis. Following peak identification, the pipeline performs comprehensive peak annotation, including genomic feature distribution across samples and peak distribution relative to transcription start sites (TSS). Enrichment quantification utilizes our custom approach detailed in the Enrichment Score Development section. The module integrates seamlessly with IGV for interactive data visualization.

### Computational Resource Management

SNAP maintains resource efficiency across various computing environments, from standard local workstations to high-performance computing clusters. Parallel processing implementation for computationally intensive steps substantially reduces processing time for large datasets. Multiple configuration options accommodate diverse computing environments, including local machines, HPC clusters, and cloud platforms (AWS). Containerized implementation using Docker/Singularity ensures reproducibility across different environments, enhancing accessibility for users with varying computational expertise and needs.

### Signal Normalization and Output Matrices

Chromatin fragments at user-defined genomic sites were quantified from BED-formatted fragment files using an R-based module integrated into SNAP, supporting both single-sample and batch analyses. Target and optional reference regions are supplied as standard BED files and imported using *rtracklayer* (v1.64.0)^49^, converted to *GenomicRanges* (v1.56.1)^50^ objects, and filtered to autosomal chromosomes by default (with optional inclusion of sex and mitochondrial chromosomes using *--include-all-chr*). Fragment overlaps with target sites are computed using *GenomicRanges::countOverlaps* to generate a site-by-sample raw count matrix. Two normalization strategies are implemented: (i) library-size normalization, which converts raw counts to counts per million (CPM), and (ii) reference-based normalization, which divides target counts by the corresponding sample’s total fragment count across user-specified reference loci (e.g., housekeeping promoters or DHS sites) to adjust for global signal-to-noise differences. All matrices, including raw, CPM-normalized, and reference-normalized, are saved in both *.RDS* and tab-delimited formats, accompanied by detailed logs summarizing input dimensions, filtering parameters, normalization settings, and output paths. Full documentation and containerized implementation are available at https://github.com/chhetribsurya/chromatin-frags-normalization.

### Output Generation with Reports

The pipeline generates key outputs that facilitate downstream analysis and interpretation. Processed BAM files with associated indices enable further analysis or integration with additional tools. Peak calls and enrichment analysis results appear in standard formats (BED, narrowPeak) compatible with existing analysis tools and databases. Fragment size distributions and coverage metrics are provided in both tabular and graphical formats, enabling straightforward result interpretation. Quality control reports and visualizations utilize *IGV-reports* (v1.12.0)^46^ and *MultiQC* (v1.25.2)^48^, providing users with clear, comprehensive analysis overviews. IGV-compatible session file generation allows rapid result loading and visualization in the IGV browser, facilitating manual review of signals at regions of interest. A detailed description of the SNAP output directories is provided in **Supplementary Table 1**.

### Datasets

CfChIP-seq datasets for H3K4me3 and H3K27ac, as well as cfMeDIP-seq and low-pass whole genome sequencing (LPWGS), were obtained from our previously published study and are accessible through GEO accession GSE243474^9^. Additional low-quality cfChIP-seq datasets were included to support the development of quality-control metrics. The details of the datasets are summarized in **Table 1**. In addition, 29 plasma samples from patients with osteosarcoma were collected and assayed by Precede Biosciences for cell-free chromatin H3K4me3, cfDNA methylation, and LPWGS (**Supplementary Table 2**). DNA methylation microarray data for 10 osteosarcoma tumor samples (GSE161407)^51^ and 35 healthy whole blood samples (GSE235717)^52^ were accessed from GEO to define hypermethylated and hypomethylated sites specific to osteosarcoma (**Supplementary Table 3**). 22 prostate cancer ChIP-seq samples from 4 individuals were accessed from GEO (GSE130408) to test for sample swaps using the SNP fingerprinting method implemented in SNAP (**Supplementary Table 4**)^53^. To identify regions of cancer-enriched H3K4me3 and H3K27ac signal, 30 ChIP-seq datasets from ENCODE and GEO were analyzed that span tumor cell lines, tissues, and white blood cells (**Supplementary Table 5**)^54,55^. 50 samples across 10 tumor types from TCGA were used for differential methylation analysis to identify regions of cancer-enriched DNA methylation (**Supplementary Table 6**)^56^. The 18-state chromHMM calls for 833 samples from the EpiMap Repository were used for H3K4me3 and H3K27ac enrichment score development^57^. 253 whole-genome bisulfite sequencing (WGBS) samples across 39 cell types and 137 donors from GSE186458 were used for MeDIP enrichment score development^58^. Cancer and healthy H3K4me3 cfChIP-seq data published by Sadeh *et al.* were downloaded from Zenodo (DOI:10.5281/zenodo.3967253) and used for external validation^10^. All data and analyses used the hg19 reference genome.

**Table 1.** Discovery Datasets.

|  | H3K4me3 | H3K27ac | MeDIP | LPWGS |
| --- | --- | --- | --- | --- |
| Total N | 601 | 340 | 298 | 239 |
| Cancer Type (N (%)) |  |  |  |  |
| Prostate adenocarcinoma | 162 (27.0%) | 62 (18.2%) | 60 (2.1%) | 26 (10.9%) |
| Neuroendocrine prostate cancer | 18 (3%) | 13 (3.8%) | 19 (6.4%) | 10 (4.2%) |
| Breast cancer | 43 (7.2%) | 37 (10.9%) | 26 (8.7%) | 15 (6.3%) |
| Colorectal cancer | 44 (7.3%) | 25 (7.4%) | 31 (10.4%) | 32 (13.4%) |
| Esophageal cancer | 15 (2.5%) | 0 (0%) | 12 (4%) | 14 (5.9%) |
| Hepatocellular cancer | 24 (4%) | 25 (7.4%) | 14 (4.7%) | 14 (5.9%) |
| Melanoma | 7 (1.2%) | 0 (0%) | 6 (2%) | 6 (2.5%) |
| Merkel cell carcinoma | 19 (3.2%) | 18 (5.3%) | 19 (6.4%) | 19 (7.9%) |
| Small-cell lung cancer | 27 (4.49%) | 15 (4.4%) | 11 (3.7%) | 7 (2.9%) |
| Non-small cell lung cancer | 110 (18.3%) | 76 (22.4%) | 33 (11.1%) | 45 (18.8%) |
| Ovarian cancer | 26 (4.3%) | 24 (7.1%) | 8 (2.7%) | 7 (2.9%) |
| Renal cancer | 12 (2%) | 9 (2.7%) | 5 (1.7%) | 2 (0.8%) |
| Small-cell bladder cancer | 2 (0.3%) | 1 (0.3%) | 2 (0.7%) | 1 (0.4%) |
| Thymic cancer | 17 (2.8%) | 16 (4.7%) | 8 (2.7%) | 9 (3.8%) |
| Glioma | 10 (1.7%) | 10 (2.9%) | 10 (3.4%) | 8 (3.3%) |
| Healthy | 64 (10.7%) | 9 (2.7%) | 34 (10.4%) | 24 (10%) |

### Plasma Processing and Cell-free Chromatin Assays for Osteosarcoma Samples

Peripheral blood from 29 unique osteosarcoma patients treated at the University of Pittsburgh Medical Center (IRB # STUDY20010034) was centrifuged at 1500 g x 10 minutes at 4 °C to obtain plasma and buffy coat on the same day of the patient encounter. Plasma Samples were immediately stored at -80 °C until experimentation. 1 mL plasma samples were shipped overnight on dry ice to Precede Biosciences, where H3K4me3 cell-free chromatin immunoprecipitation, cfDNA methylation immunoprecipitation, and LPWGS assays were performed using Precede’s automated platform. All resulting libraries were sequenced on the Illumina platform.

### Identification of Housekeeping Gene Promoter Regions

A set of 3,804 human genes with stable expression across broad cell types, defined as housekeeping genes, was obtained from Eisenberg et al^59^. Gene symbols were mapped to genomic coordinates using the *biomaRt* (v2.58.2) R package and the Ensembl *hsapiens_gene_ensembl* dataset (GRCh37/hg19 assembly)^60^. For each gene, the promoter region was defined as the 2,000 base pairs upstream of the transcription start site. Genes located on autosomes (chromosomes 1–22) were retained, and coordinates were formatted in BED format for downstream analyses, resulting in 3,402 housekeeping promoter regions (**Supplementary Table 7**).

### Identification of H3K4me3 and H3K27ac Cancer-Enriched Regions

Peak BED files for H3K4me3 and H3K27ac were obtained from ENCODE/GEO for multiple tumor-derived cell lines and tissues, along with primary immune-cell references (**Supplementary Table 5**). BED files were imported into R as GRanges objects using the *GenomicRanges* package (v1.56.1). For each histone mark, tumor/tissue peak sets were intersected across all available cancer sources to define a common pan-cancer peak core. A unified WBC reference was constructed by merging immune-cell peak sets, and overlapping regions were removed from the pan-cancer core to limit to sites with low signal from chromatin blood cell-derived chromatin. The remaining peaks were restricted to autosomes (chr1–chr22), yielding final sets of WBC-depleted pan-cancer regions with 416 H3K4me3 peaks (**Supplementary Table 8**) and 43 H3K27ac peaks (**Supplementary Table 9**). These curated peak sets were used for cfChIP-seq quantification and classification analyses.

### Identification of Cancer-Enriched Hypermethylated Regions

DNA methylation β-values were obtained from Illumina EPIC arrays for whole blood from GSE235717 (**Supplementary Table 3**) using the *SeSAMe* (v1.22.1) R package^61^. Probe-level filtering was performed using Illumina EPIC manifests to exclude non-CpG, sex chromosome, and generally masked probes^62,63^. Tumor methylation data from ten TCGA cancer types (**Supplementary Table 6**) were subsetted to the same probe set and merged with the WBC controls, yielding a combined dataset of 85 samples, including 35 WBC and 50 tumor samples. Differentially methylated probes were identified using the *limma* (v3.60.3), which applies linear modeling with empirical Bayes moderation of standard errors to improve variance estimation and statistical power^64^. For each comparison, probes with adjusted p-values < 0.05 and a methylation difference ≥0.5 were selected as significantly hypermethylated. These CpG sites were mapped to genomic coordinates using the Illumina EPIC hg19 annotation. To define methylated regions, we extended each CpG site by ±500 bp and merged overlapping intervals using the *IRanges* (v2.38.0) package. The resulting 1,051 non-overlapping regions were exported as BED files for downstream cfMeDIP-seq signal quantification (**Supplementary Table 10**).

### Identification of Osteosarcoma Hyper- and Hypo-methylated Regions

DNA methylation data for osteosarcoma and whole blood from healthy donors were obtained from the GEO under accession numbers GSE161407 and GSE235717, respectively^51,52^. Raw IDAT files were processed using the *SeSAMe* (v1.22.1) R package for quality control, normalization, and extraction of methylation β-values^61^. Probes with detection p-values > 0.01, those located on sex chromosomes, and known cross-reactive or SNP-affected probes were removed^62,63^. Differential methylation analysis was performed using the *limma* (v3.60.3) package^64^. A linear modeling framework was used to identify differentially methylated positions (DMPs) between osteosarcoma and whole blood samples. The empirical Bayes moderation approach implemented in eBayes was applied to stabilize variance estimates and improve the detection of DMPs^64^. CpG sites with an absolute methylation difference > 0.3 and FDR < 0.01 were considered significantly differentially methylated. Sites with increased methylation in osteosarcoma were designated as hypermethylated, and those with decreased methylation were designated as hypomethylated. To define genomic regions surrounding these loci, ±500 base pair windows were added around the coordinates of each differentially methylated CpG site, and the resulting regions were exported in BED format for downstream analyses, resulting in 1,188 hypermethylated regions and 2,983 hypomethylated regions for osteosarcoma compared to whole blood (**Supplementary Table 11**).

### Enrichment Score Development

To quantify the effectiveness of each targeted assay in enriching for the targeted epigenetic marks, we developed an enrichment score based on read density in defined on- and off-target regions. For cfChIP-seq marks, we utilized the 18-state chromHMM segmentation of the hg19 genome across 833 biosamples from the Epimap resource^57^. The genome was partitioned into 200 bp intervals, each annotated with one of the 18 chromHMM states, resulting in a matrix of 200 bp genomic windows by 833 samples. On-target regions for H3K4me3 and H3K27ac were selected based on chromHMM states corresponding to the respective histone marks. Specifically, 200 bp windows were retained as on-target sites if >95% of Epimap samples annotated them with a chromHMM state characteristic of the respective mark. For H3K27ac, on-target states included 1_TssA, 3_TssFlnkU, 8_EnhG2, and 9_EnhA1; for H3K4me3, on-target states included 1_TssA, 2_TssFlnk, 3_TssFlnkU, 4_TssFlnkD, 8_EnhG2, and 14_TssBiv. We further removed windows overlapping hg19 exclusion regions or located on chromosomes X and Y. The resulting BED files were sorted, merged, and filtered to retain only regions larger than 1000 bp. Off-target regions were defined using a complementary approach: windows that were never annotated with the mark-specific chromHMM states in any sample. Exclusion regions and chromosomes X and Y were again removed, and regions within 10 kb of any on-target site were also removed. After sorting, merging, and filtering for size (>1000 bp), final on- and off-target region sets were established. For H3K4me3, the final on-target and off-target lists contained 8,174 and 281,426 regions, respectively. For H3K27ac, the final on-target and off-target lists contained 1,651 and 219,809 regions, respectively. For cfMeDIP-seq, on-target regions were defined as CpG loci with methylation beta values >0.90 across 253 samples from the Human Methylome Atlas^58^, excluding chromosomes X and Y. These CpGs were mapped to 1000 bp genomic windows to form the on-target set (n = 1,186). Off-target regions were defined as CpG loci with methylation beta values <0.10, mapped in the same manner, using a 1:1 ratio to the on-target number of sites, yielding 1146 off-target regions. Complete lists of on- and off-target regions can be found in **Supplementary Table 12**. For each assay and sample, we calculated the total base pairs in all on-target (on_bp) and off-target (off_bp) regions, as well as the number of fragments overlapping these regions (on_fragments, off_fragments). The enrichment score was then computed as the ratio of normalized read densities in on- versus off-target regions:

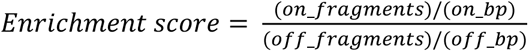

This metric provides a robust, quantitative measure of enrichment by immunoprecipitation for cfChIP-seq (H3K4me3, H3K27ac) and cfDNA methylation experiments.

### Comparison of ChIP-seq Quality Metrics Between Plasma and Tissue Samples

To evaluate how quality metrics differ between tissue-derived and circulating chromatin samples, we compared fragment count, enrichment score, and peak number between our plasma cfChIP-seq data and publicly available ENCODE ChIP-seq datasets from tissues and cell lines^54^. For this comparison, we selected a representative panel of H3K4me3 and H3K27ac ChIP-seq datasets from ENCODE (listed in **Supplementary Table 13**), spanning multiple cell types, including HeLa, HCT116, HepG2, K562, and primary immune cells. For ENCODE samples, BAM files were used to calculate fragment counts and enrichment scores, following the same procedure applied to plasma data. Peak numbers were extracted directly from the corresponding narrow peak BED files. Comparisons were performed separately for H3K4me3 and H3K27ac assays using violin plots, and statistical significance was assessed using t tests.

### Quality Metrics Optimization Methods

Enrichment score cut-offs were optimized using a knee-point algorithm^65^. For each assay and region, classification performance for distinguishing cancer from healthy samples was evaluated across a grid of enrichment score thresholds. At each candidate threshold, samples were subset, and ROC/AUC values were calculated using the *pROC* (v1.18.5) package. Thresholds with insufficient observations, missing classes, or constant predictors were skipped. The AUC–threshold curve was smoothed with a centered moving average, normalized, and compared against the straight line connecting its endpoints. Local maxima of the distance curve were identified as candidate knees, and the first that satisfied an upward-turn criterion (increase in slope, increase in mean level, or position above the secant between neighbors) was chosen, provided that the unsmoothed value was not lower than the baseline at the minimum threshold. The x-coordinate of this point was reported as the optimized enrichment score cut-off. One optimized cut-off was obtained per region and then summarized across regions to define assay-specific distributions. Fragment count cut-offs were optimized using a parallel elbow-point algorithm. Instead of classification performance, signal stability was assessed by computing the coefficient of variation (CV) across Eisenberg housekeeping gene promoter regions at increasing fragment count thresholds (50,000 increments). CV–threshold curves were processed in the same way as above, except that candidate elbows were required to satisfy a downward-turn criterion (reduced slope, reduced mean level, or concavity below the local secant) and to yield a CV lower than the baseline. The selected elbow x-coordinate was reported as the optimized fragment count cut-off, and distributions across regions were summarized for each assay.

### ctDNA Estimation

ichorCNA is a widely used tool for estimating ctDNA fractions from plasma, based on tumor-associated copy number alterations in LPWGS data^66^. However, ichorCNA is not suitable for cell-free ChIP-seq and MeDIP-seq data due to the coverage biases of these data types. To address this, the SNAP pipeline incorporates Fragle, a method that estimates ctDNA levels based on the density distribution of cfDNA fragment lengths^34^. We used Fragle to estimate ctDNA content directly from cfChIP and cfMeDIP data, and compared these values to estimates obtained using ichorCNA on LPWGS data from the same plasma samples. For each assay, Fragle ctDNA values were calculated using all genomic sites, as well as stratified by on-target and off-target regions. For H3K4me3 and H3K27ac, on- and off-target sites matched those used for enrichment score calculation. For MeDIP, on-target sites were defined similarly, while off-target sites encompassed regions not included as on-target sites due to the limited number of available loci.

### Osteosarcoma Differential Epigenomic Analyses

Plasma cfDNA from 29 patients with osteosarcoma was profiled for cell-free chromatin H3K4me3, cfDNA methylation, and LPWGS, and all data were processed through SNAP. Direct outputs from SNAP were used for the analyses. For differential analysis, samples were stratified by the ctDNA fraction estimated using Fragle^34^. The five samples with the highest ctDNA fractions were defined as the high-ctDNA group, and the five with the lowest fractions as the low-ctDNA group. Fragment counts were quantified by overlapping mapped reads with peak regions to generate raw count matrices. Differential peak analysis between high and low ctDNA groups was performed using *DESeq2*^67^ (v1.46.0) with median-of-ratios normalization to adjust for sequencing depth. The pipeline included size factor and dispersion estimation, fitting of negative binomial models, and Wald testing. Peaks with p < 0.05 were considered significant, with positive and negative log₂ fold-change values denoting up- and downregulation, respectively. Significant peaks were annotated with the nearest gene using *GENCODE v19*^68^, defining promoter regions as ±2 kb from transcription start sites. Gene ontology (GO) enrichment was conducted with *clusterProfiler*^69^ (v4.14.6) and *org.Hs.eg.db*, converting gene symbols to Entrez IDs via *bitr()*. Separate enrichment tests for up- and downregulated peaks were performed across biological process (BP), molecular function (MF), and cellular component (CC) domains using hypergeometric testing with Benjamini–Hochberg correction (adjusted p < 0.05). Only GO terms containing 10–500 genes were retained for interpretation.

### SNAP Resource Usage Benchmark

To assess SNAP’s computational performance and resource efficiency, we benchmarked the pipeline using the osteosarcoma cohort comprising 58 plasma samples profiled with IP-based epigenomic assays. All analyses were performed on a workstation equipped with an AMD Ryzen Threadripper PRO 5975WX processor (32 physical cores, 64 logical cores) and 503 GB RAM, running Ubuntu 24.04.3 LTS. Subsets of increasing size (10, 20, 30, 40, 50, and 58 samples) were randomly selected from the full cohort and processed independently using SNAP with identical parameters. For each run, total wall-clock runtime and computational resource usage were recorded to evaluate scalability and efficiency as a function of cohort size.

All statistical analyses and visualizations were performed in R (v4.4.0) using the methods and parameters detailed above.

## Results

### Difference in QC Metrics between Cell-free and Tissue ChIP-seq

We observed stark differences in quality metrics between plasma cfChIP-seq and ENCODE tissue ChIP-seq datasets. For both H3K4me3 and H3K27ac assays, tissue-derived samples showed significantly higher peak numbers (**Supplementary Figure 1A**, H3K4me3: p = 3.5e-05; H3K27ac: p = 2.1e-05) and fragment counts (**Supplementary Figure 1B**, H3K4me3: p = 4.8e-05; H3K27ac: p = 8.3e-04) compared to plasma samples. Previous ENCODE and related studies have proposed that high-quality H3K4me3 ChIP-seq datasets should contain ≥20 million unique fragments and ≥20,000 narrow peaks^31,32,70^. However, these thresholds are not practical for H3K4me3 cfChIP-seq, as most samples would be classified as low-quality under such criteria. Similarly, for H3K27ac ChIP-seq, quality has been benchmarked at ≥20 million fragments and ≥50,000 narrow peaks^31,32,71^, yet these standards are also not directly applicable to H3K27ac cfChIP-seq (**Supplementary Figure 1**, red dashed lines indicate tissue ChIP-seq quality cut-offs). In contrast, enrichment scores, which measure signal over background at target-specific genomic regions, showed minimal differences between sample types (**Supplementary Figure 1C**, H3K4me3: p = 0.89; H3K27ac: p = 0.03). These results demonstrate that traditional QC cutoffs designed for tissue or cell-line derived ChIP-seq data are poorly suited for evaluating cfChIP-seq data, given the lower and more variable signal inherent to circulating chromatin.

### Quality Metrics Establishment

Peak number is often used as a basic indicator of ChIP-seq data quality. However, in cfChIP-seq datasets, relying solely on peak number can be misleading. Visualization of signal intensity tracks revealed that samples with high peak counts but low enrichment or fragment counts exhibited highly variable and noisy signals (**Supplementary Figure 2**). To establish more reliable measures, we evaluated two complementary quantitative metrics. Fragment count reflects the total number of isolated cfDNA fragments, while enrichment score quantifies how effectively those fragments are enriched at expected regulatory regions. Across assays, these two metrics were largely uncorrelated, suggesting that they capture distinct aspects of experimental quality (**Supplementary Figure 3**). Fragment count primarily reflects input quantity and sequencing depth, whereas enrichment score measures immunoprecipitation efficiency. Together, these findings demonstrate that peak number alone is insufficient and support the use of enrichment score and fragment count as complementary, orthogonal metrics for assessing cfChIP-seq and cfMeDIP-seq data quality.

Having selected fragment count and enrichment score as key quality parameters, we next sought to define empirical cut-offs for these measures. We reasoned that data of sufficient quality should capture biological signals that enable accurate performance in classification tasks. For simplicity, we focused on the task of distinguishing the presence or absence of cancer-derived chromatin. We performed a classification task comparing cancer samples with >10% ctDNA against healthy samples, using individual regions with cancer-enriched H3K4me3 (**Supplementary Table 8**), H3K27ac (**Supplementary Table 9**), and hypermethylated regions (**Supplementary Table 10**). For each region, classification performance was evaluated across a series of enrichment score thresholds in 0.5-unit increments. A knee-point algorithm (see Methods) was then applied to select the optimized cut-off for each region. Examples of region-level optimization are shown in **Supplementary Figure 4A**, and the distributions of optimized enrichment score cut-offs across all regions are presented in **Figure 2A**. For H3K4me3, cut-offs centered around a mean of 8.7, while H3K27ac and MeDIP showed means of 8.5 and 2.9, respectively. These results demonstrate that excluding samples with low enrichment scores improves the ability to identify biologically relevant chromatin signatures in cfChIP and cfMeDIP data. Based on these findings, we propose a general enrichment score threshold of approximately 9 for histone mark assays and 3 for MeDIP, acknowledging that optimal values may vary depending on assay type, sequencing depth, and specific analytical objectives.

**Figure 2.**
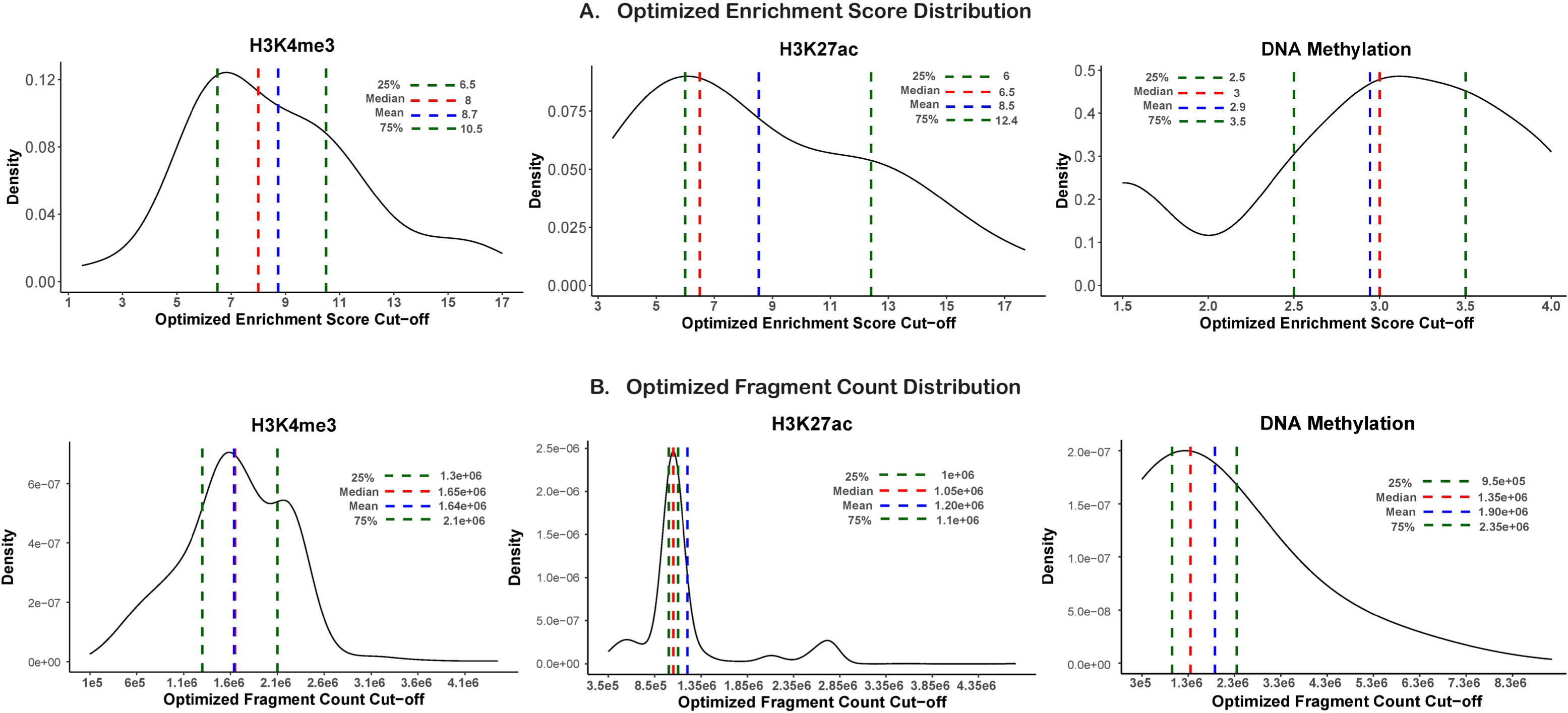
Optimized quality control thresholds for enrichment score and fragment count across cfChIP-seq and cfMeDIP-seq assays. **(A)** Distribution of optimized enrichment score cut-offs obtained using individual regions for H3K4me3, H3K27ac, and DNA methylation. Cut-offs were derived using a classification task distinguishing plasma samples with >10% ctDNA from healthy controls across region-specific enrichment thresholds. The optimal value for each region was determined using a knee-point algorithm. Dashed vertical lines denote the 25^th^-75^th^ percentile (green), median (red), and mean (blue). **(B)** Distribution of optimized fragment count cut-offs for H3K4me3, H3K27ac, and DNA methylation data. Thresholds were determined by minimizing signal variability (coefficient of variation) across Eisenberg housekeeping gene promoter regions, with optimal values selected using an elbow-point algorithm. Dashed vertical lines represent the 25^th^-75^th^ percentile (green), median (red), and mean (blue).

Unlike enrichment score, which reflects specific capture of the targeted epitope and can therefore be optimized using classification performance, fragment count primarily reflects whether sufficient DNA was captured to generate stable signals. When fragment counts are low, signals become more variable due to stochasticity. To identify optimal fragment count thresholds, we set the objective of minimizing signal variability in the Eisenberg housekeeping gene promoter regions. We performed cut-off optimization based on the coefficient of variation (CV) of the signal in the regions. For each region, CV was evaluated across a series of fragment count thresholds in 50,000-unit increments. An elbow-point algorithm (see Methods) was then applied to select the optimized cut-off for each region. Examples of region-level optimization are shown in **Supplementary Figure 4B**, and the distributions of optimized fragment count cut-offs across all regions are presented in **Figure 2B**. For H3K4me3, fragment count cut-offs centered around a mean of 1,640,000, while H3K27ac and MeDIP showed means of 1,200,000 and 1,900,000, respectively. These results indicate that samples with very low fragment counts yield unstable or noisy signals, and filtering such samples improves data consistency and biological interpretability. Based on these findings, we recommend a general fragment count threshold of approximately 1,000,000 – 2,000,000 fragments, with the understanding that optimal values may vary depending on assay type and sequencing depth.

### Quality Metrics Validation

To evaluate the robustness of our quality control thresholds, we applied them to an independent cfChIP-seq dataset of H3K4me3 from cancer and healthy samples published by Sadeh et al^10^. Samples were stratified into high-quality (N = 164) and low-quality (N = 66) groups based on mean optimized cut-offs for enrichment score (> 8.7) and fragment count (> 1,640,000). Using 416 cancer-enriched H3K4me3 regions (**Supplementary Table 8**), application of these filters consistently improved classification performance, as reflected by higher AUC values (**Supplementary Figure 5A**), and reduced variability in 3,402 housekeeping regions, as indicated by lower CV (**Supplementary Figure 5B**). Together, these results confirm our approach for identifying quality thresholds and show that enrichment score and fragment count provide robust and generalizable quality control metrics for IP-based epigenomic profiling of circulating chromatin.

### Assessment of Enrichment Score Specificity Across Assays

To evaluate the specificity of the enrichment score metric, we calculated scores across all samples and compared distributions by antibody type. Each antibody yielded high enrichment scores for its corresponding mark, with minimal signal in non-target assays except for H3K4me3 and H3K27ac, as they both mark active promoters (**Supplementary Figure 6**). These results confirm the specificity and interpretability of the enrichment score metric and support its utility as a robust sample-level quality measure within the SNAP pipeline.

### Fragle ctDNA estimation

Fragle ctDNA estimates showed strong concordance with ichorCNA when applied to LPWGS data, consistent with the benchmarking context reported in its original publication^34^ (Pearson’s r = 0.78, p < 2.2e-308, RMSE = 13%; **Supplementary Figure 7**). When extended to cfChIP-seq data, using off-target sites for both H3K4me3 and H3K27ac, correlations remained high though slightly attenuated (H3K4me3: Pearson’s r = 0.71, p < 2.2e-308, RMSE = 16%; H3K27ac: Pearson’s r = 0.66, p < 2.2e-308, RMSE = 16%) (**Supplementary Figure 8**). For comparison, MeDIP-seq achieved the strongest correlations overall when evaluated on both genome-wide and off-target sites (r = 0.72, p < 2.2e-308, RMSE = 15%, **Supplementary Figure 8**). To assess the ability of each method to distinguish cancer from healthy plasma, we computed area under the ROC curve (AUC) values. Fragle-based ctDNA estimates from H3K4me3, H3K27ac, and LPWGS data slightly outperformed ichorCNA-derived estimates from LPWGS. Specifically, Fragle achieved AUCs of 0.88 for H3K4me3, 0.87 for H3K27ac, 0.84 for LPWGS, and 0.73 for MeDIP, compared with an AUC of 0.82 applying ichorCNA to LPWGS data (**Supplementary Figure 9A**). Optimal cut-offs for cancer and healthy classification were determined using Youden’s index (Fragle: H3K4me3 = 7%, H3K27ac = 5%, MeDIP = 10%, LPWGS = 5%; ichorCNA LPWGS = 2%; **Supplementary Figure 9B-C**). These cut-offs indicate that although Fragle generally shows higher sensitivity, its effective limit of detection is marker-dependent, with distinct thresholds required for H3K4me3, H3K27ac, and MeDIP. These findings demonstrate that Fragle can be applied to cfChIP-seq and cfMeDIP-seq data, achieving classification performance comparable to ichorCNA.

### *SMaSH* SNP Fingerprint Sample Clustering

We tested the ability of *SMaSH* to identify sample mismatches based on SNP fingerprints derived from IP-based data. We first applied SMaSH to public ChIP-seq datasets where multiple marks were profiled from the same samples. Analyzing 22 prostate cancer ChIP-seq samples from 4 individuals in the GEO dataset GSE130408^53^, *SMaSH* showed that most ChIP-seq profiles from the same individual clustered together. However, it detected a swap where FOXA1 ChIP-seq profiles from consecutively numbered samples were likely switched with each other (**Supplementary Figure 10**). We then extended this evaluation to cfDNA data by analyzing 20 samples from 5 individuals, each profiled with cfChIP-seq (H3K4me3 and H3K27ac), cfMeDIP-seq, and LPWGS. In this setting, *SMaSH* achieved perfect clustering by individual, demonstrating its robustness for sample matching across multiple cfDNA assay types (**Supplementary Figure 11**). Based on these results, *SMaSH* has been fully integrated into the SNAP pipeline to enable detection of potential sample mismatches.

### Osteosarcoma Plasma Epigenomic Profiling using SNAP

To evaluate the performance of SNAP on external datasets and across additional cancer types, we applied the pipeline to plasma samples from patients with osteosarcoma, a rare cancer type not included in the initial training cohorts. We analyzed 29 plasma samples from patients with clinically diagnosed osteosarcoma using H3K4me3, cfDNA methylation, and LPWGS assays. All data were processed through the SNAP pipeline. Tumor fraction (ctDNA) was estimated from ichorCNA on LPWGS data and Fragle on H3K4me3 and cfDNA methylation data, revealing positive correlations between ichorCNA and Fragle estimated ctDNA (**Supplementary Figure 12**). Genomic annotation of the peaks revealed that H3K4me3 peaks were predominantly enriched at promoters (**Supplementary Figure 13A**), as expected, while cfDNA methylation peaks were enriched in distal intergenic and intronic regions (**Supplementary Figure 13B**). Quality control metrics, enrichment score, and total fragment count were evaluated for all samples (**Supplementary Figure 14**).

In the cfDNA methylation data, signal intensity in regions previously identified as hypermethylated in osteosarcoma relative to white blood cells showed a strong positive correlation with ctDNA levels (Pearson’s r = 0.92, p = 1.5e-12, **Figure 3A**), while signal in hypomethylated regions exhibited a significant negative correlation (Pearson’s r = -0.68, p = 4.5e-05, **Figure 3B**). Additionally, the ratio of hyper- to hypomethylation signals demonstrated the highest correlation with ctDNA fraction (Pearson’s r = 0.99, p = 8.2e-16, **Figure 3C**).

**Figure 3.**
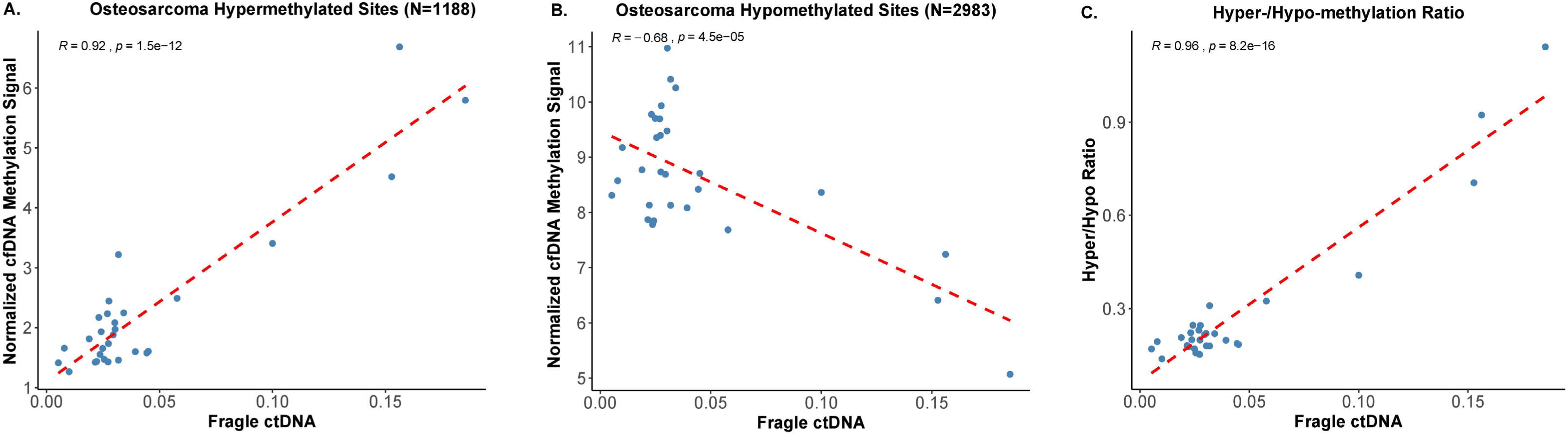
Correlation between osteosarcoma-enriched cfDNA methylation signals and ctDNA fraction. **(A)** Normalized cfDNA methylation signal across 1,188 regions identified as hypermethylated in osteosarcoma compared to white blood cells, showing a strong positive correlation with Fragle-estimated ctDNA (Pearson’s r = 0.92, p = 1.5e-12). **(B)** Normalized cfDNA methylation signal across 2,983 hypomethylated regions exhibiting a significant negative correlation with ctDNA levels (Pearson’s r = –0.68, p = 4.5e-05). **(C)** The ratio of hyper- to hypomethylation signals demonstrates the strongest positive correlation with ctDNA fraction (Pearson’s r = 0.99, p = 8.2e-16). Each point represents an individual plasma sample (n = 29), and red dashed lines indicate linear regression fits.

Differential H3K4me3 peak analysis in high-versus low-ctDNA samples (Fragle ctDNA top 5 vs bottom 5) at transcription start sites (TSSs) identified 550 upregulated and 545 downregulated peaks (p < 0.05; **Figure 4A**). Notably, we observed upregulation of several genes with established roles in osteosarcoma pathogenesis, including WISP1 (log₂FC =1.69, p = 5.9e-06), which enhances VEGF-A expression and promotes angiogenesis^72^, BMP-7 (log₂FC = 0.96, p = 0.02), a bone morphogenetic protein highly expressed in osteosarcoma and involved in osteoblast differentiation^73,74^, LRRC15 (log₂FC =1.87, p = 6.2e-05), previously shown to be overexpressed in osteosarcoma tissues^75,76^, and TGFB2 (log₂FC = 0.65, p = 5.1e-04), a key regulator of epithelial-mesenchymal transition and metastasis in late-stage cancers^77,78^ (**Figure 4A**). Gene ontology analysis of the upregulated H3K4me3 peaks revealed substantial enrichment for developmental pathways (**Figure 4B**). Among the top 20 enriched biological processes, were gene sets expected to be active in osteosarcoma, including bone-specific developmental pathways, with ossification (adjusted p = 2.1e-06), embryonic skeletal system development (adjusted p = 6.3e-06), skeletal system morphogenesis (adjusted p = 1.2e-05), and osteoblast differentiation (adjusted p = 1.6e-05; **Figure 4B**).

**Figure 4.**
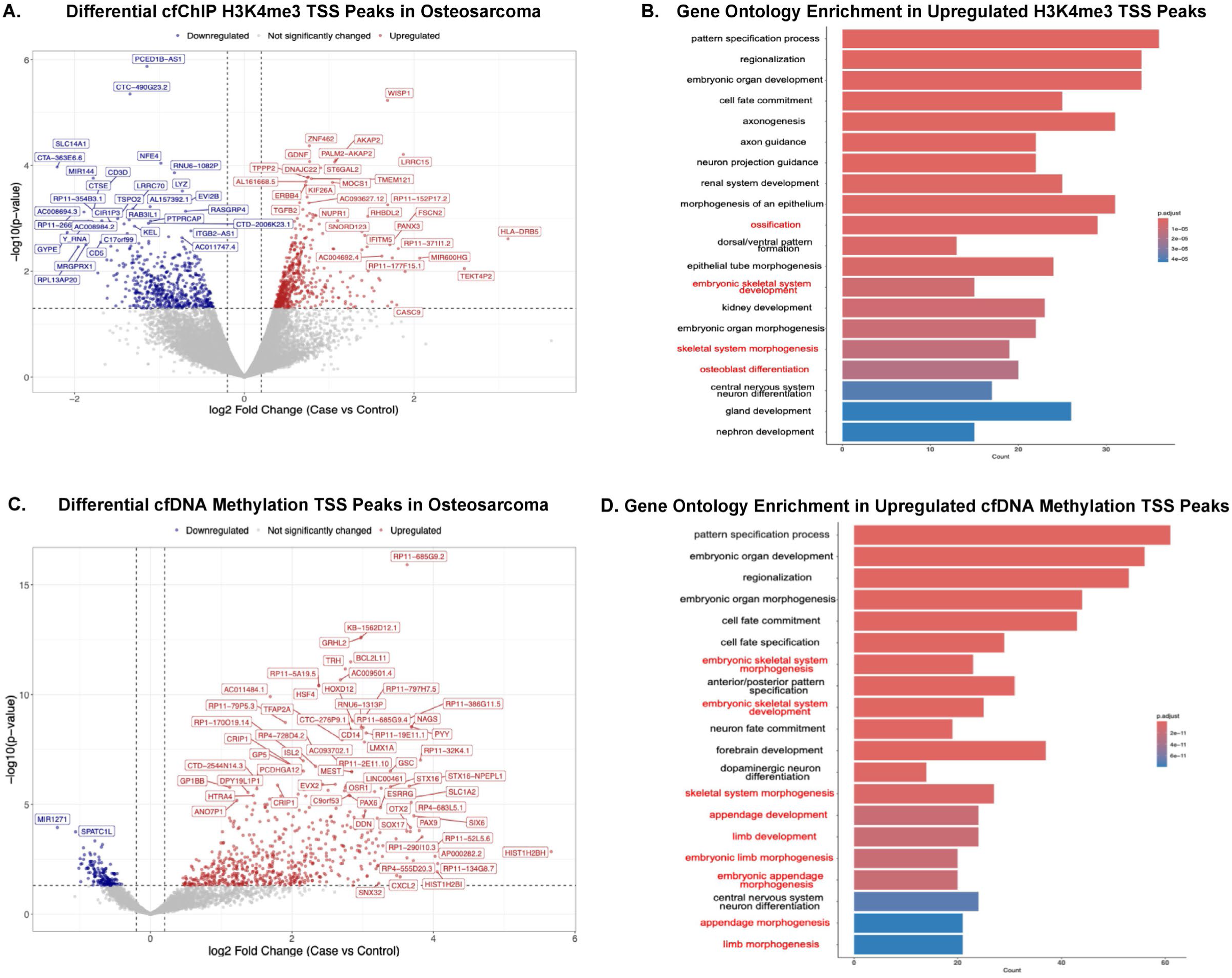
Differential epigenomic analysis of osteosarcoma plasma samples stratified by Fragle-estimated ctDNA fraction. **(A)** Volcano plot showing differential H3K4me3 peaks at transcription start sites (TSS ± 2 kb) between high- and low-ctDNA osteosarcoma samples. Red and blue dots indicate significantly up- and downregulated peaks, respectively (p < 0.05), and gray dots denote non-significant changes. Representative upregulated osteosarcoma-associated genes are labeled. **(B)** Gene ontology over-representation analysis of genes linked to differential H3K4me3 peaks. Bars represent –log₁₀(adjusted p values), with skeletal, limb, and bone development pathways highlighted in red. **(C)** Volcano plot showing differential DNA methylation at TSS regions from cfDNA methylation analysis between high- and low-ctDNA groups. **(D)** GO enrichment analysis of differentially methylated regions (DMRs) from cfDNA methylation, showing significant enrichment for developmental and morphogenesis pathways (highlighted in red).

Differential cfDNA methylation TSS peak analyses revealed 635 hypermethylated and 161 hypomethylated regions in high versus low ctDNA samples (**Figure 4C**). The hypermethylated regions included promoters of key transcription factors: TFAP2A (log₂FC = 2.71, p = 1.1e-08), a recently identified driver of osteosarcoma metastasis^79,80^, PAX6 (log₂FC = 2.82, p = 4e-06), associated with key developmental roles and poor prognosis in osteosarcoma and other cancers^81–84^, and CRIP1 (log₂FC = 2.04, p = 1.9e-07), which shows complex expression patterns in osteosarcoma with potential prognostic value^85^ (**Figure 4C**). Gene enrichment analysis of hypermethylated regions demonstrated strong enrichment for skeletal and limb developmental processes (**Figure 4D**), with embryonic skeletal system morphogenesis (adjusted p = 4.3e-17), embryonic skeletal system development (adjusted p = 7.2e-16), skeletal system morphogenesis (adjusted p = 4.7e-12), and multiple limb/appendage development pathways (adjusted p = 1.6e-11 to 8e-11) among the most significantly enriched (**Figure 4D**).

SNP fingerprinting within SNAP confirmed that samples from both H3K4me3 and cfDNA methylation (**Supplementary Figure 15**) assays originated from the same individuals. In summary, these results demonstrate how SNAP functions as a robust data processing pipeline that generates high-quality outputs that enable the detection of cancer-enriched epigenomic alterations in plasma.

### Computational Performance and Scalability of SNAP

To evaluate the computational efficiency and scalability of SNAP, we assessed resource usage using the osteosarcoma cohort with increasing sample sizes (10, 20, 30, 40, 50, and 58 samples). Total wall-clock runtime increased in a near-linear manner with cohort size, rising from approximately 8 hours for 10 samples to 45 hours for the full 58-sample dataset (**Supplementary Figure 16**). Across all runs, SNAP was executed using a fixed allocation of 16 CPU cores, while memory requirements scaled moderately with input size, increasing from 128 GB for smaller subsets to a maximum of 256 GB for cohorts of ≥50 samples. Raw input data size ranged from 35.3 GB (10 samples) to 251.1 GB (58 samples). These results demonstrate that SNAP scales predictably with cohort size and can efficiently process dozens of IP-based cfDNA samples on a single high-memory workstation without requiring changes to core pipeline configuration.

## Discussion

Characterizing epigenomic features of cancer from circulating chromatin is becoming a critical focus in precision oncology. Liquid biopsy approaches that profile circulating chromatin histone modifications and DNA methylation hold promise for early cancer detection, longitudinal monitoring, and therapeutic stratification^8,9,86,87^. However, the inherently low input, high fragmentation, variable quality of circulating chromatin, and pre-analytical variability introduced during sample collection and processing, present unique challenges that limit the effectiveness of traditional ChIP-seq and MeDIP-seq bioinformatics pipelines^28–30^.

In this study, we developed and validated SNAP, a reproducible and scalable workflow specifically designed for cfDNA epigenomic profiling using cfChIP-seq and cfMeDIP-seq data. Building on recent advances in cfDNA fragmentomics and immunoprecipitation-based methods^9,88^, SNAP incorporates quality control metrics optimized for cfDNA data, a novel enrichment scoring system to assess assay performance, and ctDNA estimation based on fragment length distributions. Importantly, SNAP is implemented as a modular and parameterized workflow, enabling straightforward extension to additional histone modifications or epigenomic assays by updating assay-specific configuration files rather than core pipeline logic. Our benchmarking results demonstrate that applying SNAP’s QC framework improves cancer classification performance across multiple assays and cancer types, including H3K4me3 ChIP-seq, H3K27ac ChIP-seq, and MeDIP-seq. SNAP further provides standardized, figure-ready visualizations and summary outputs for QC metrics, enrichment scores, and ctDNA estimates, facilitating rapid data interpretation and cross-assay comparison. The integration of *SMaSH*-based SNP fingerprinting further enhances sample integrity by enabling accurate identity verification and cross-assay linkage, minimizing the risk of sample mix-ups in multi-assay workflows^35^. This capability is particularly useful as epigenomic liquid biopsy approaches are enabling longitudinal profiling of the same individuals and parallel analysis of multiple chromatin marks from the same sample. Additionally, incorporating Fragle-based ctDNA estimation allows direct quantification of tumor fraction from cfChIP-seq and cfMeDIP-seq data, supporting downstream applications requiring ctDNA estimates^34^. SNAP is distributed with well-documented defaults and user-adjustable parameters, allowing investigators to tailor QC thresholds and analysis settings to specific assays, sample types, and study designs.

In an independent osteosarcoma plasma cohort, SNAP identified tumor-specific H3K4me3 and 5mC signatures that closely tracked with ctDNA abundance and reflected key gene regulatory programs associated with sarcoma biology. The consistent enrichment of skeletal morphogenesis, ossification, and limb developmental pathways across assays highlights the robustness of cfDNA-based epigenomic profiling for capturing disease-associated signatures. As a highly aggressive bone malignancy with limited measurable disease on imaging, osteosarcoma remains difficult to monitor using conventional tools^89–92^. To our knowledge, this represents the first application of cell-free chromatic epigenomic profiling of osteosarcoma, demonstrating that SNAP can detect tumor-derived epigenomic signals from plasma and enable minimally invasive assessment of gene regulatory programs in this disease.

While our study presents SNAP as a reproducible and scalable workflow tailored for cfDNA epigenomic profiling, several limitations should be acknowledged. First, Fragle-based ctDNA estimation in SNAP relies on fragmentomic features and exhibits marker-dependent sensitivity. Although we demonstrated broad concordance between Fragle and ichorCNA estimates, particularly at moderate to high ctDNA fractions, variability increased at lower tumor fractions and differed across chromatin marks. The effective limit of detection for Fragle was higher than that of ichorCNA and varied by assay (H3K4me3 ≈7%, H3K27ac ≈5%, MeDIP ≈10%). These findings indicate that Fragle estimates below mark-specific thresholds should be interpreted with caution. In settings where ctDNA levels are expected to be very low, orthogonal approaches with higher analytical sensitivity, such as targeted hybrid capture or digital PCR, may be more appropriate for precise tumor fraction estimation. In addition, fragmentomic signatures are influenced by biological context. While Fragle produced reliable ctDNA estimates in plasma, performance in other liquid biopsy sources, such as urine or CSF, remains uncertain. Users of SNAP should therefore be aware that ctDNA estimation accuracy may vary depending on assay type and sample context. Second, SNAP was validated using H3K4me3 cfChIP-seq, H3K27ac cfChIP-seq, and cfMeDIP-seq data, but its QC metrics and enrichment scoring framework have not yet been evaluated on other circulating epigenomic marks that also hold promise as proxies of transcriptional activity in liquid biopsy studies. These include gene body–associated active marks such as H3K36me3 and 5-hydroxymethylcytosine (5hmC), promoter- and enhancer-associated active marks such as H3K4me2, and constitutive heterochromatin–associated repressive marks such as H3K9me3. Systematic validation across these assays will be important to establish the generalizability of SNAP to epigenomic features with distinct regulatory roles.

In summary, SNAP addresses a critical gap in circulating chromatin epigenomics by providing a reproducible, flexible, and biologically informed analysis framework tailored to the unique properties of cell-free histone modification and DNA methylation immunoprecipitation data. As liquid biopsy applications continue to expand within oncology and in non-oncologic disease, SNAP will support the use of cfChIP-seq and cfMeDIP-seq for studying disease biology and eventually guiding treatment decisions. The pipeline’s modular architecture also makes it amenable to support additional cfDNA-based assays, e.g., H3K36me3, H3K4me2, H3K9me3, and 5hmC^93^, facilitating broader integration into future workflows for biologic discovery and biomarker development.

## Acknowledgements

*Biorender.com* was used for the Graphic Abstract and Figure 1. Osteosarcoma biospecimen collection was facilitated as part of the Musculoskeletal Oncology Tumor Registry (MOTOR), which is funded, in part, by Pittsburgh Cure Sarcoma. The MOTOR works with the Pitt Biospecimen Core at the University of Pittsburgh. Work performed in the Pitt Biospecimen Core (RRID:SCR_025229) and services and instruments used in this project were supported, in part, by the University of Pittsburgh, the Office of the Senior Vice Chancellor for Health Sciences. We thank the patients who generously donated samples to the biobank, without whom this research would not have been possible.

## Author contributions

Ze Zhang (Data curation [equal], Investigation [equal], Software [supporting], Visualization [equal], Writing—original draft [lead], Writing—review & editing [equal]); Paulo Da Silva Cordeiro (Data curation [equal], Investigation [equal], Software [lead], Visualization [equal], Writing—original draft [equal], Writing—review & editing [equal]); Surya B. Chhetri (Data curation [equal], Investigation [equal], Software [equal], Visualization [equal], Writing—original draft [equal], Writing—review & editing [equal]); Brad Fortunato ([Data curation [equal], Investigation [equal], Software [equal], Writing—review & editing [equal]); Razane El Hajj Chehade [Data curation [equal], Investigation [equal], Software [supporting], Writing—review & editing [equal]); Karl Semaan ([Data curation [equal], Investigation [equal], Software [supporting], Writing—review & editing [equal]); Gunsagar Gulati ([Data curation [equal], Investigation [equal], Software [supporting], Writing—review & editing [equal]); Garyoung Gary Lee ([Data curation [equal], Investigation [equal], Software [supporting], Writing—review & editing [equal]); Chris Hemauer ([Data curation [equal], Investigation [equal], Software [supporting], Writing—review & editing [equal]); Zhenjie Jin ([Data curation [equal], Investigation [equal], Writing—review & editing [equal]); Weiwei Bian ([Data curation [equal], Investigation [equal], Writing—review & editing [equal]); Shahabeddin Sotudian ([Data curation [equal], Investigation [equal], Writing—review & editing [equal]); Ziwei Zhang ([Data curation [equal], Investigation [equal], Writing—review & editing [equal]); David Osei-Hwedieh ([Data curation [equal], Investigation [equal], Writing—review & editing [equal]); Corrie Painter ([Data curation [equal], Writing—review & editing [equal]); Tanya E. Heim ([Data curation [equal], Writing—review & editing [equal]); Kurt R. Weiss ([Data curation [equal], Writing—review & editing [equal]); Rashad Nawfal ([Data curation [equal], Writing—review & editing [equal]); Marc Eid ([Data curation [equal], Writing—review & editing [equal]); Damien Vasseur ([Data curation [equal], Writing—review & editing [equal]); John Canniff ([Data curation [equal], Writing—review & editing [equal]); Hunter Savignano ([Data curation [equal], Writing—review & editing [equal]); Noa Phillips ([Data curation [equal], Writing—review & editing [equal]); Ji-Heui Seo ([Data curation [equal], Project administration [supporting], Supervision [supporting], Writing—review & editing [equal]); Matthew L. Freedman ([Data curation [equal], Project administration [supporting], Supervision [supporting], Resources [supporting], Writing—review & editing [equal]);, Sylvan C. Baca ([Data curation [equal], Project administration [lead], Supervision [lead], Resources [lead], Writing—original draft [supporting], Writing—review & editing [equal]).

## Supplementary data

Supplementary Figures 1–16 are provided in the file Supplementary_Figures.doc.

Supplementary Tables 1–13 are available in the file Supplementary_Tables.xlsx.

## Data and code availability

The SNAP pipeline is publicly available at https://github.com/prc992/SNAP/.

Publicly available datasets used in this study were obtained from the EpiMap Repository (https://compbio.mit.edu/epimap/), TCGA (https://portal.gdc.cancer.gov/), GTEx (https://www.gtexportal.org/home/), and GEO (https://www.ncbi.nlm.nih.gov/geo/) under the accession numbers GSE220160, GSE256092, GSE243474, GSE161407, GSE235717, GSE130408, GSE161948, GSE120738, and GSE186458.

Due to privacy restrictions, osteosarcoma plasma sequencing data are available from the corresponding author upon reasonable request and execution of a data use agreement.

## Author Notes

Ze Zhang, Paulo Da Silva Cordeiro, and Surya Chhetri should be regarded as co-first authors.

