## Supplementary Figures 1-16 for "SNAP: Streamlined Nextflow Analysis Pipeline for Immunoprecipitation-Based Epigenomic Profiling of Circulating Chromatin"

Table of Contents

**Supplementary Figure 1.** Comparison of quality control metrics between plasma cfChIP-seq and ENCODE tissue ChIP-seq datasets for **A.** peak number (the red dashed line represents the commonly used quality control peak number cut-off for tissue ChIP-seq: H3K4me3: > 20,000; H3K27ac: > 50,000); **B.** fragment count (the red dashed line represents the commonly used quality control fragment count cut-off for tissue ChIP-seq: H3K4me3: > 2e+07; H3K27ac: 2e+07); **C.** enrichment score.	3

**Supplementary Figure 2.** cfChIP-seq H3K4me3 IGV track comparisons across high-quality and low-quality samples.	4

**Supplementary Figure 3.** Correlation comparisons across fragment count, enrichment score, and peak number in **A.** H3K4me3 cfChIP-seq, **B.** H3K27ac cfChIP-seq, and **C.** cfMeDIP assays.	5

**Supplementary Figure 4. A.** Optimized individual region enrichment score cut-off selection using a knee-point algorithm; **B.** Optimized individual region fragment count cut-off selection using an elbow-point algorithm	6

**Supplementary Figure 5.** Validation of optimized quality control thresholds using an independent cfChIP-seq dataset. **A**. Classification performance of cancer versus healthy samples from the *Sadeh et al.^1^* cfChIP-seq H3K4me3 dataset dichotomized by applying SNAP-defined quality control thresholds (enrichment score > 8.7 and fragment count > 1.64e+06); **B.** Coefficient of variation (CV) of normalized signal intensities across Eisenberg housekeeping gene promoters in all samples from the *Sadeh et al.^1^* cfChIP-seq H3K4me3 data dichotomized by applying SNAP-defined quality control thresholds	7

**Supplementary Figure 6.** H3K4me3, H3K27ac, and MeDIP enrichment score comparison across cell-free H3K4me3, H3K27ac, MeDIP, and LPWGS assays	8

**Supplementary Figure 7.** Correlation between LPWGS ichorCNA ctDNA and Fragle ctDNA	9

**Supplementary Figure 8.** Correlations between ichorCNA ctDNA and Fragle ctDNA across cfChIP-seq H3K4me3, cfChIP-seq H3K27ac, and cfMeDIP-seq assays	10

**Supplementary Figure 9.** **A.** ROC curves for using ctDNA to distinguish cancer plasma from healthy plasma samples; **B.** Fragle ctDNA comparison between cancer plasma and healthy plasma samples across H3K4me3 cfChIP-seq, H3K27ac cfChIP-seq, and cfMeDIP assays; **C.** LPWGS Fragle ctDNA and ichorCNA ctDNA comparisons between cancer plasma and healthy plasma samples.	11

**Supplementary Figure 10.** *SMaSH* SNAP fingerprint clustering identifies mismatched ChIP-seq samples from GSE130408^2^	12

**Supplementary Figure 11.** *SMaSH* SNP fingerprinting for MeDIP, LPWGS, H3K4me3, and H3K27ac samples from the same individuals	13

**Supplementary Figure 12. A.** Osteosarcoma plasma samples: Correlation between LPWGS ichorCNA-estimated ctDNA and H3K4me3 Fragle-estimated ctDNA; **B.** Correlation between LPWGS ichorCNA-estimated ctDNA and cfDNA methylation Fragle-estimated ctDNA	14

**Supplementary Figure 13.** Osteosarcoma plasma H3K4me3 cfChIP-seq and cfDNA methylation peak signal distribution in **A.** genomic locations and **B.** transcription start site locations	15

**Supplementary Figure 14.** Quality control metrics for osteosarcoma plasma H3K4me3 cfChIP-seq and cfDNA methylation samples. Red dashed lines represent established quality control cut-offs (cell-free chromatin H3K4me3: enrichment score > 8.7 and fragment count > 1.64e+06; cfDNA methylation: enrichment score > 2.9 and fragment count > 1.2e+06)	16

**Supplementary Figure 15.** *SMaSH* SNP fingerprinting for osteosarcoma plasma H3K4me3 cfChIP-seq and cfDNA methylation samples	17

**Supplementary Figure 16.** Computational performance and resource usage of SNAP on the osteosarcoma cohort	18

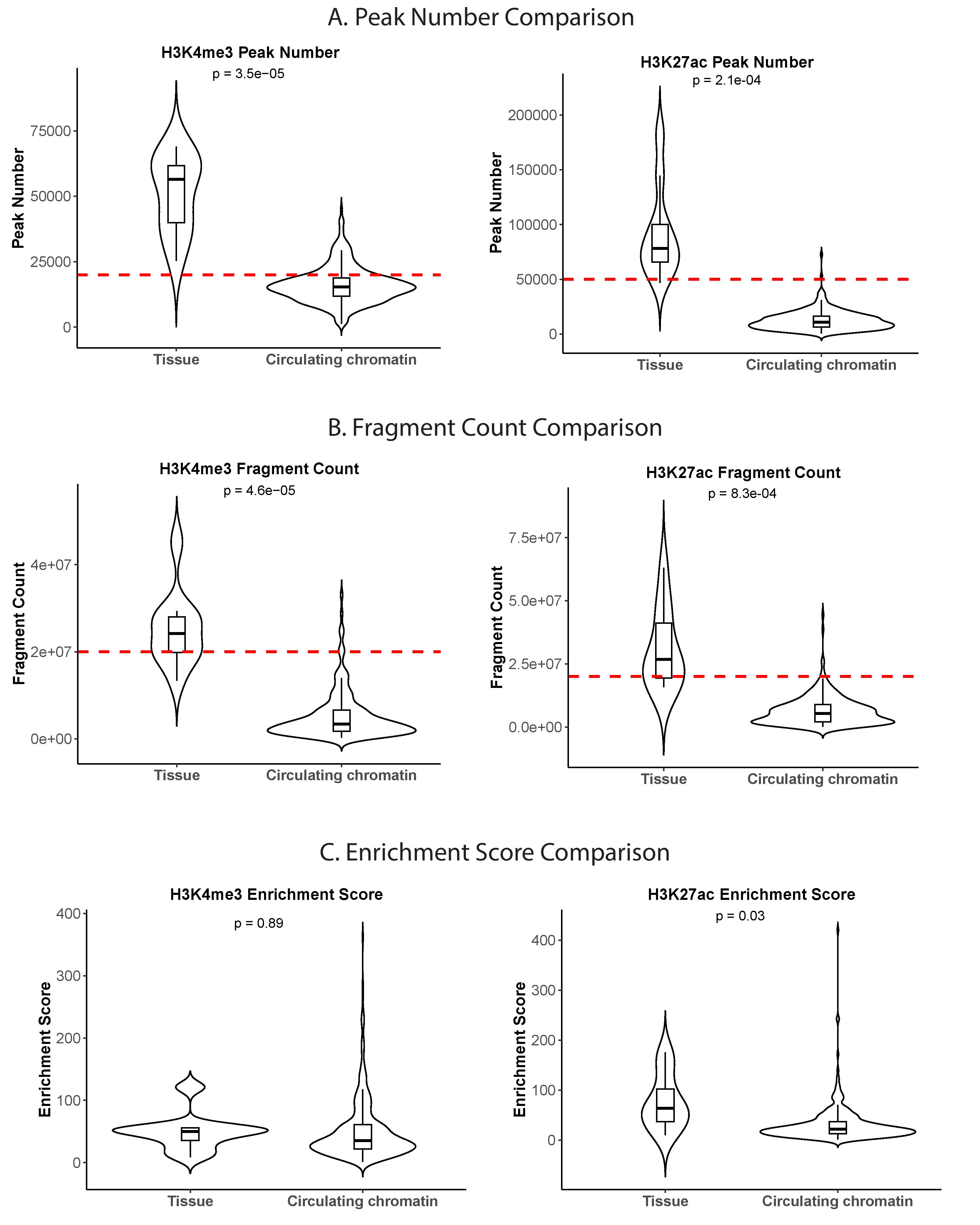

**Supplementary Figure 1.** Comparison of quality control metrics between plasma cfChIP-seq and ENCODE tissue ChIP-seq datasets for **A.** peak number (the red dashed line represents the commonly used quality control peak number cut-off for tissue ChIP-seq: H3K4me3: > 20,000; H3K27ac: > 50,000); **B.** fragment count (the red dashed line represents the commonly used quality control fragment count cut-off for tissue ChIP-seq: H3K4me3: > 2e+07; H3K27ac: 2e+07); **C.** enrichment score. *Note that ENCODE tissue H3K4me3 datasets were generated using different antibodies than plasma cfChIP-seq (e.g., Abcam ab8580), which has been reported to exhibit cross-reactivity with H3K4me2 and may contribute to higher apparent peak numbers in tissue samples. Nonetheless, plasma cfChIP-seq shows consistently lower signal yield and complexity than tissue ChIP-seq across all metrics.*

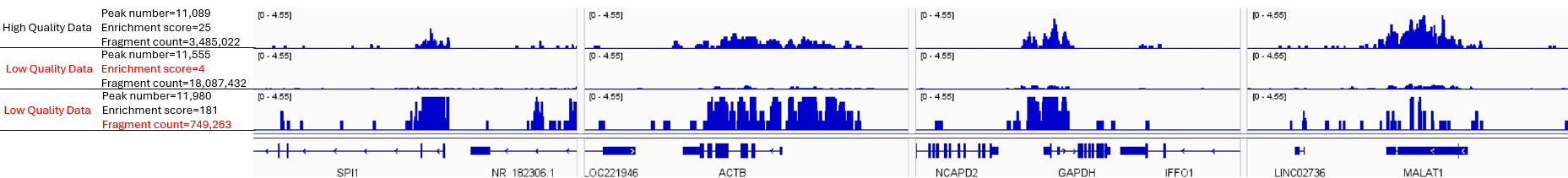

**Supplementary Figure 2.** cfChIP-seq H3K4me3 IGV track comparisons across high-quality and low-quality samples.

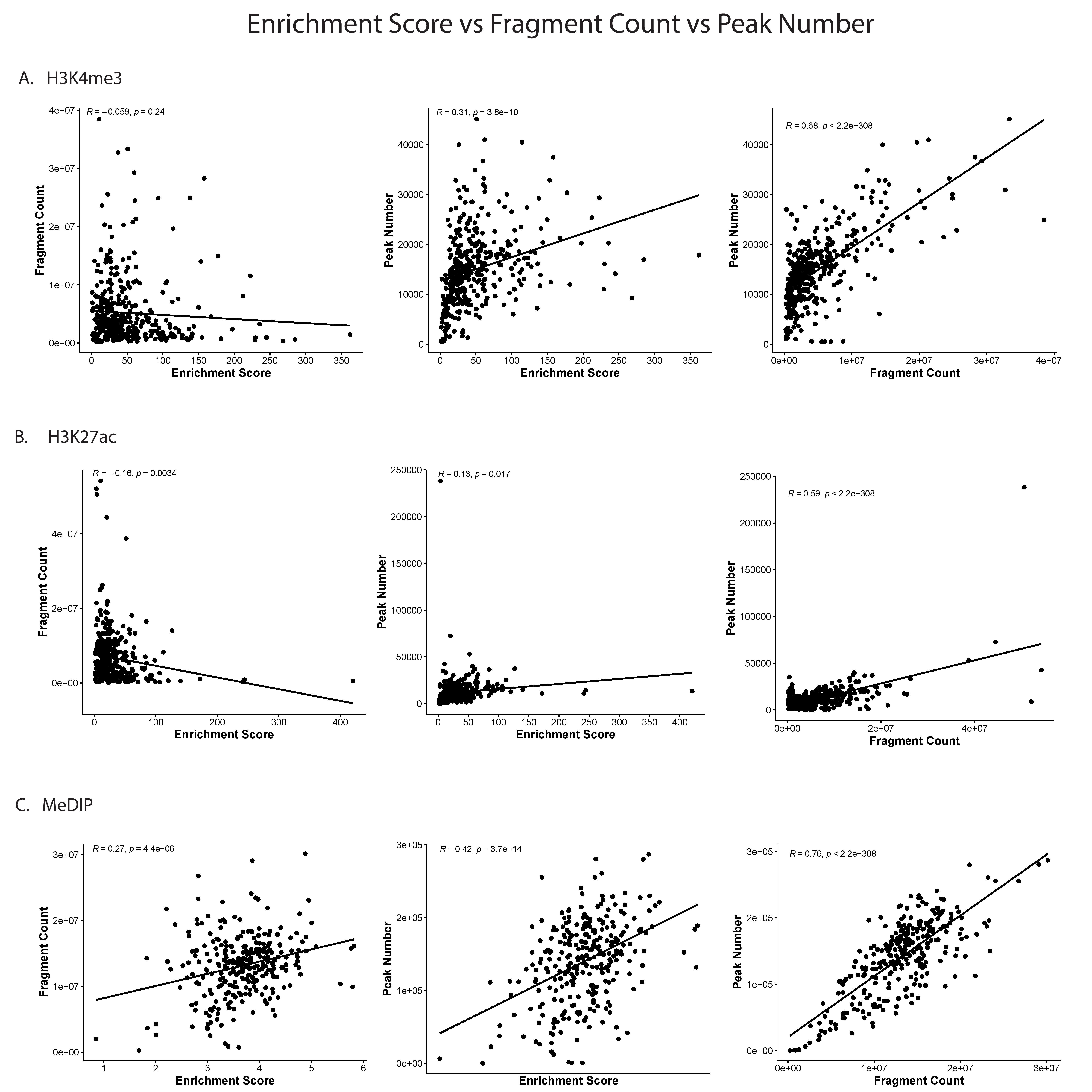

**Supplementary Figure 3.** Correlation comparisons across fragment count, enrichment score, and peak number in **A.** H3K4me3 cfChIP-seq, **B.** H3K27ac cfChIP-seq, and **C.** cfMeDIP assays.

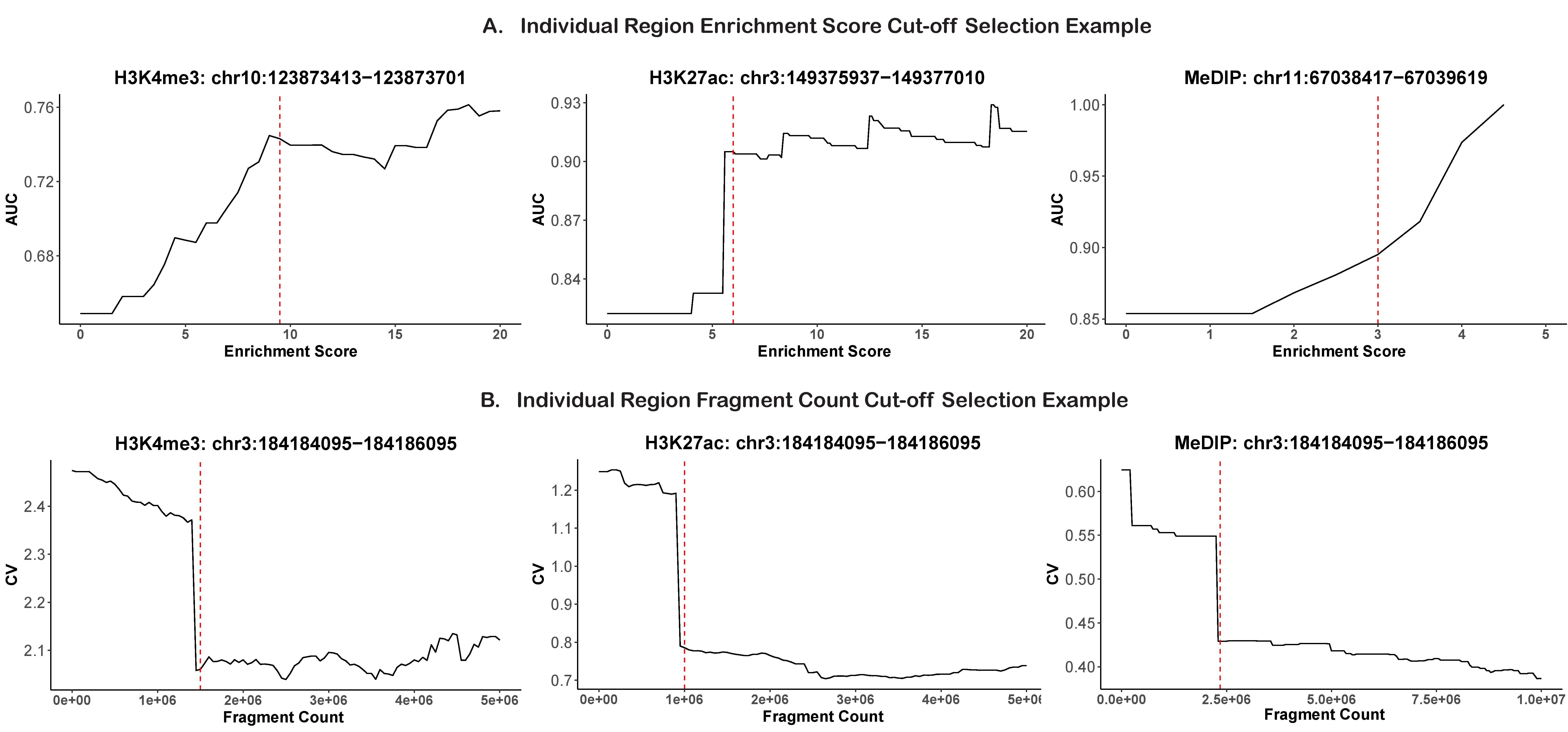

**Supplementary Figure 4. A.** Optimized individual region enrichment score cut-off selection using a knee-point algorithm; **B.** Optimized individual region fragment count cut-off selection using an elbow-point algorithm.

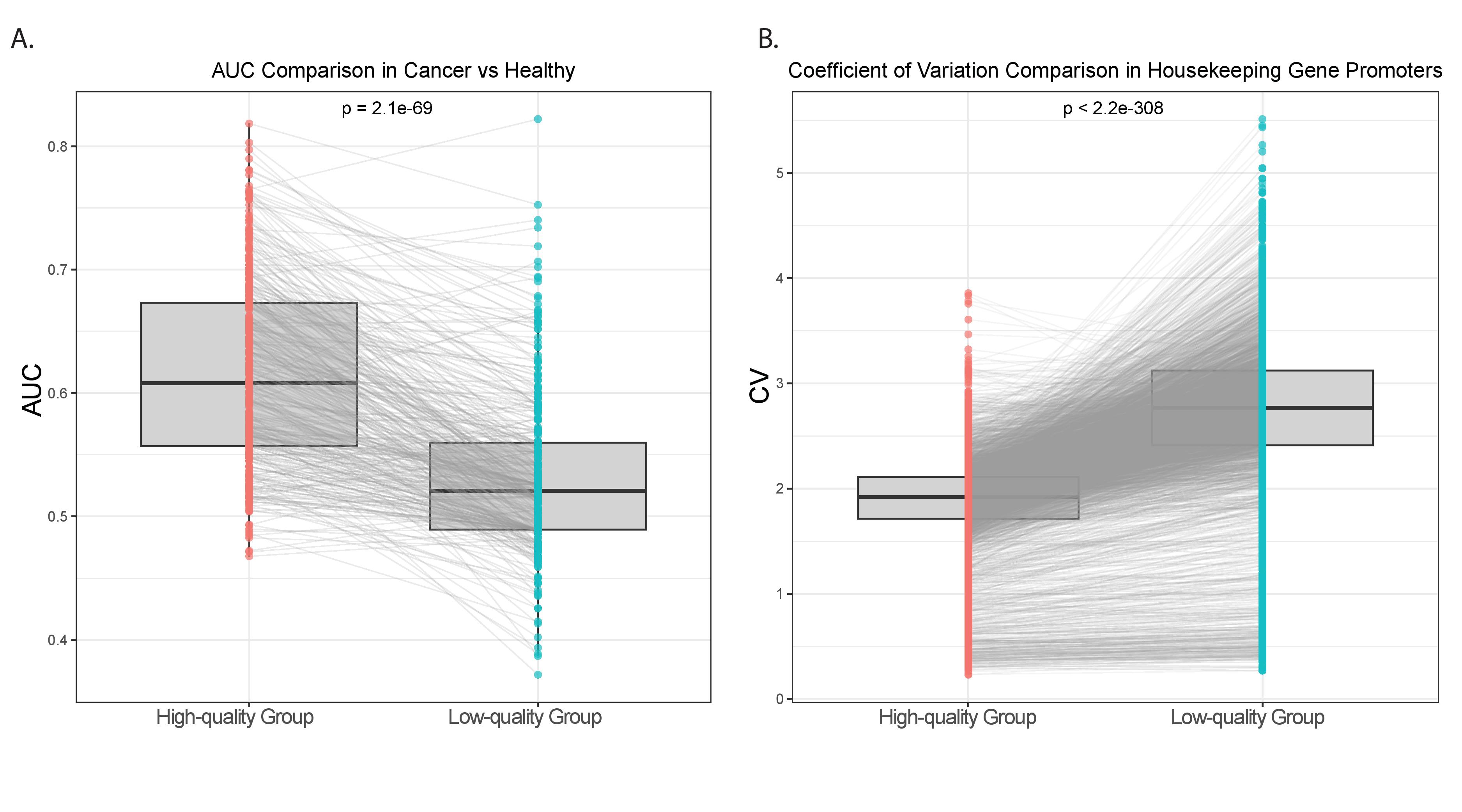

**Supplementary Figure 5.** Validation of optimized quality control thresholds using an independent cfChIP-seq dataset. **A**. Classification performance of cancer versus healthy samples from the *Sadeh et al.^1^* cfChIP-seq H3K4me3 dataset, using 416 cancer-enriched H3K4me3 regions, dichotomized by applying SNAP-defined quality control thresholds (enrichment score > 8.7 and fragment count > 1.64e+06); **B.** Coefficient of variation (CV) of normalized signal intensities across 3,402 Eisenberg housekeeping gene promoters in all samples from the *Sadeh et al.^1^* cfChIP-seq H3K4me3 data dichotomized by applying SNAP-defined quality control thresholds.

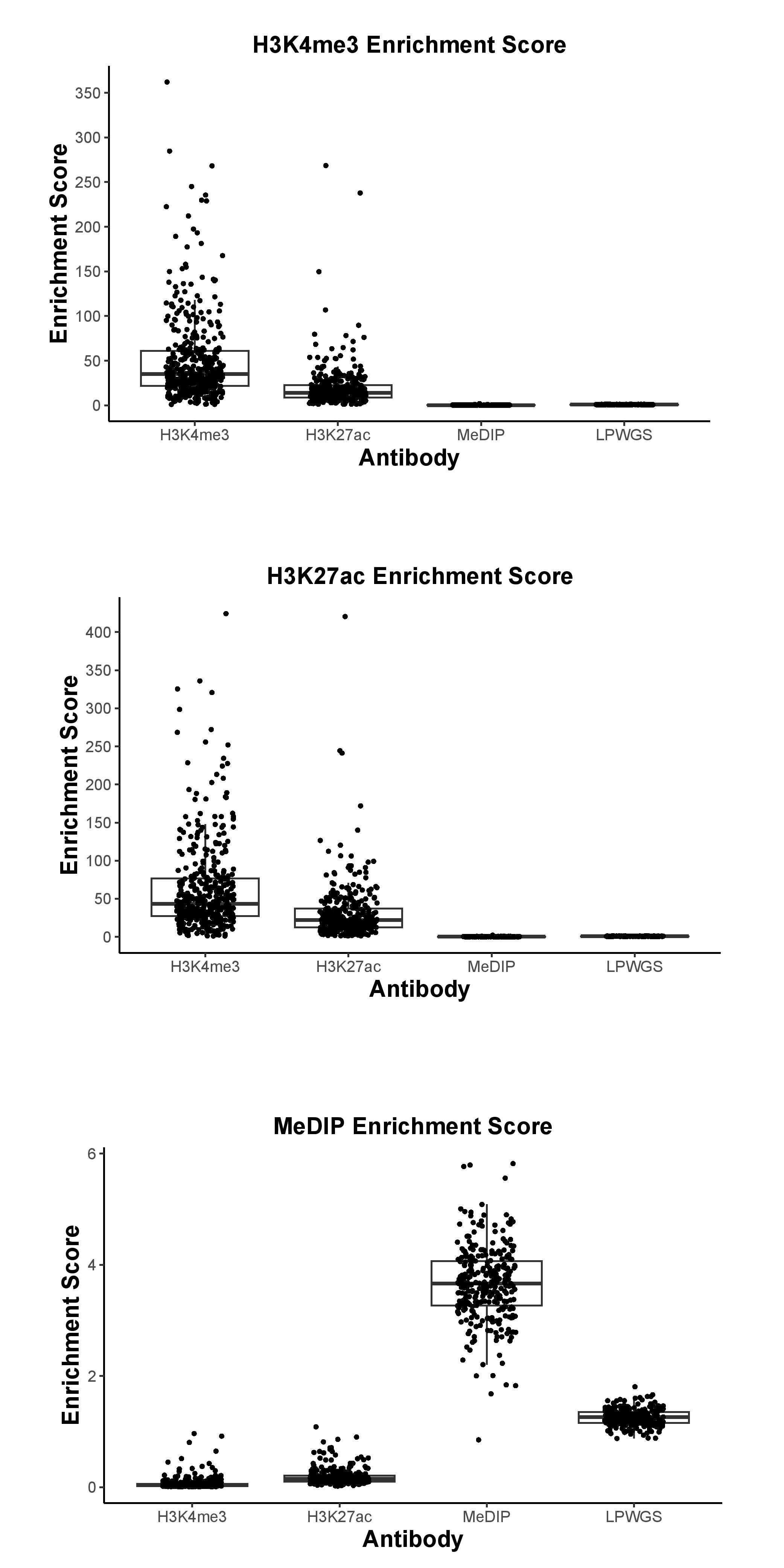

**Supplementary Figure 6.** H3K4me3, H3K27ac, and MeDIP enrichment score comparison across cell-free H3K4me3, H3K27ac, MeDIP, and LPWGS assays.

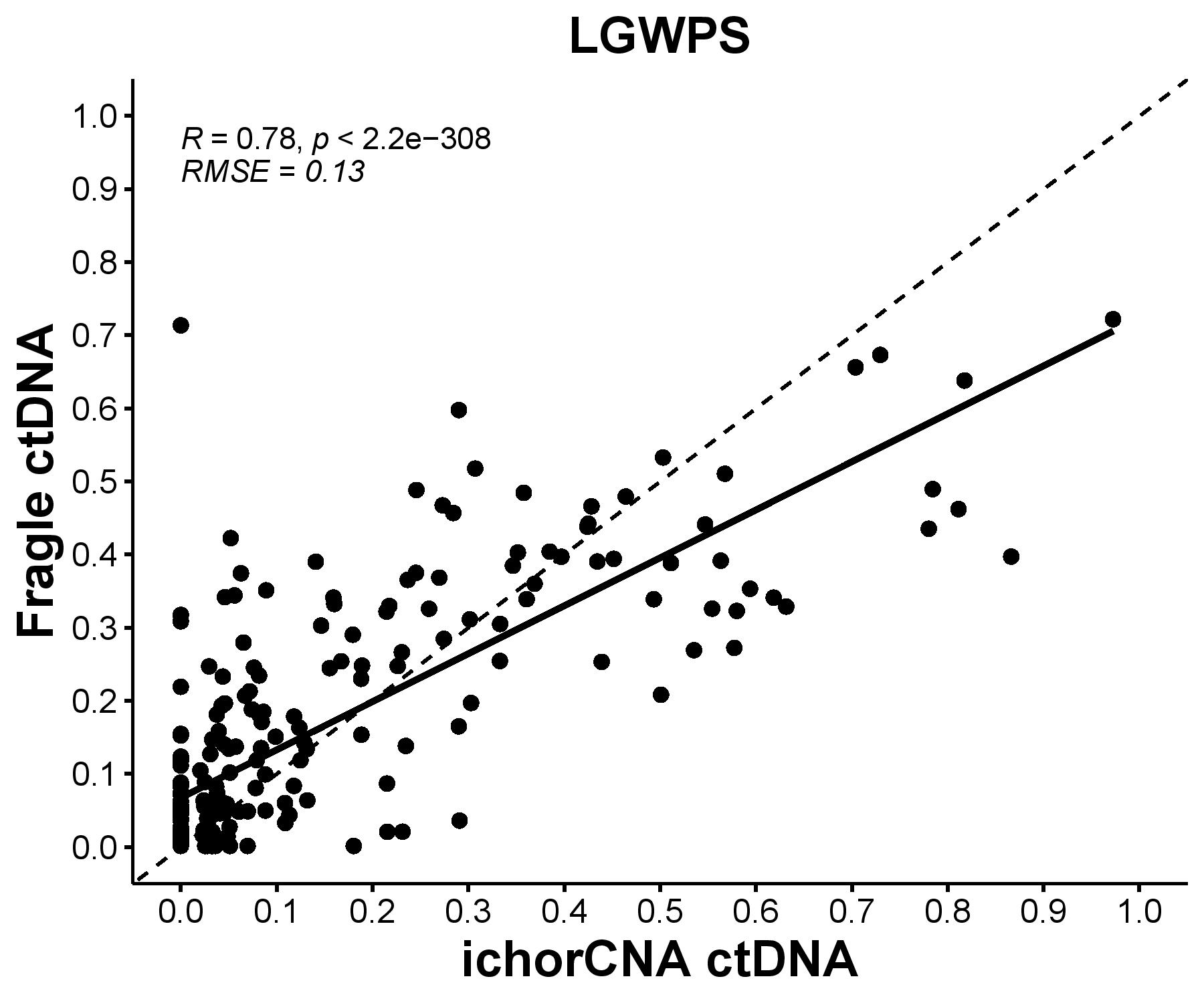

**Supplementary Figure 7.** Correlation between LPWGS ichorCNA ctDNA and Fragle ctDNA.

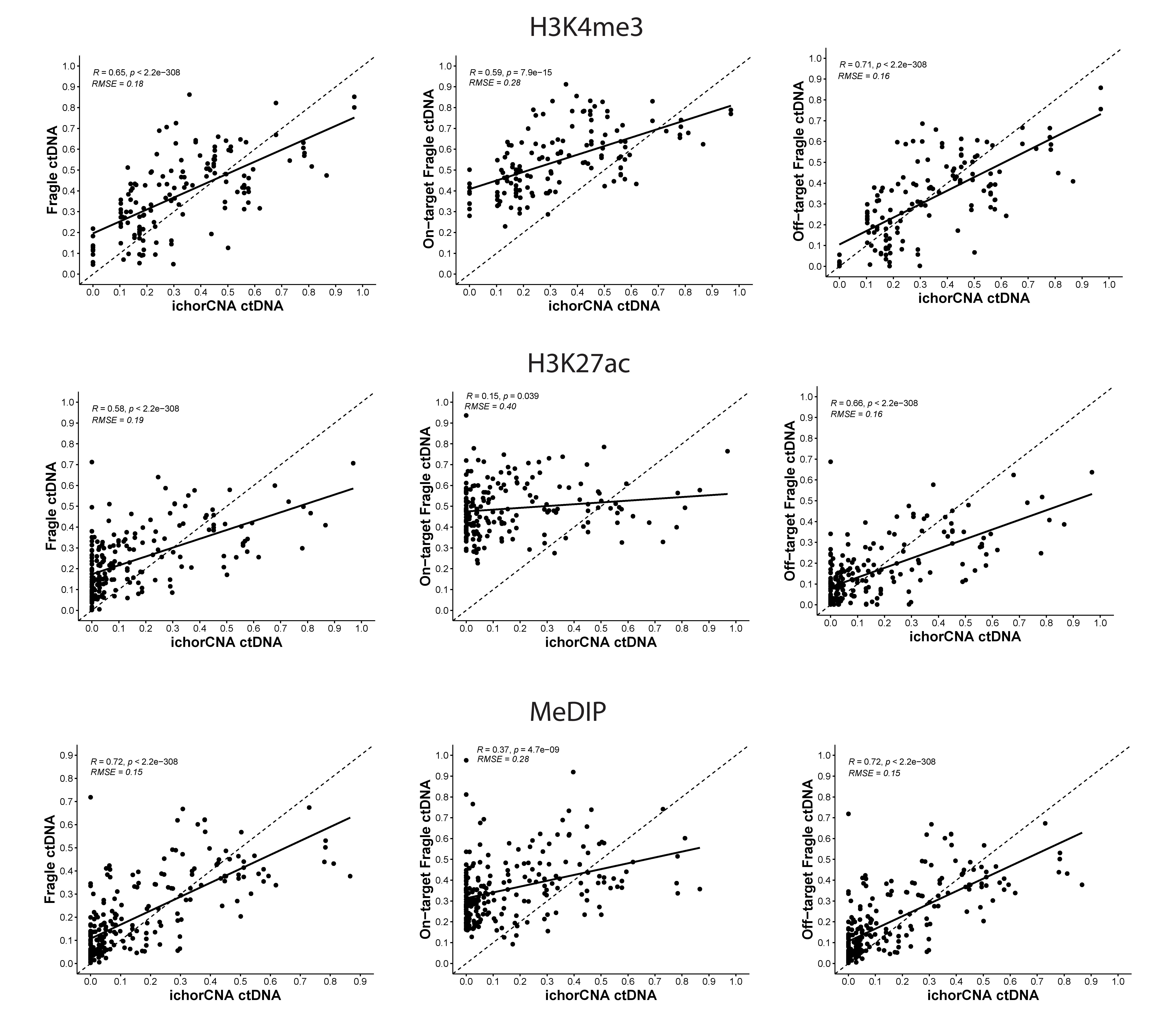

**Supplementary Figure 8.** Correlations between ichorCNA ctDNA and Fragle ctDNA across cfChIP-seq H3K4me3, cfChIP-seq H3K27ac, and cfMeDIP-seq assays.

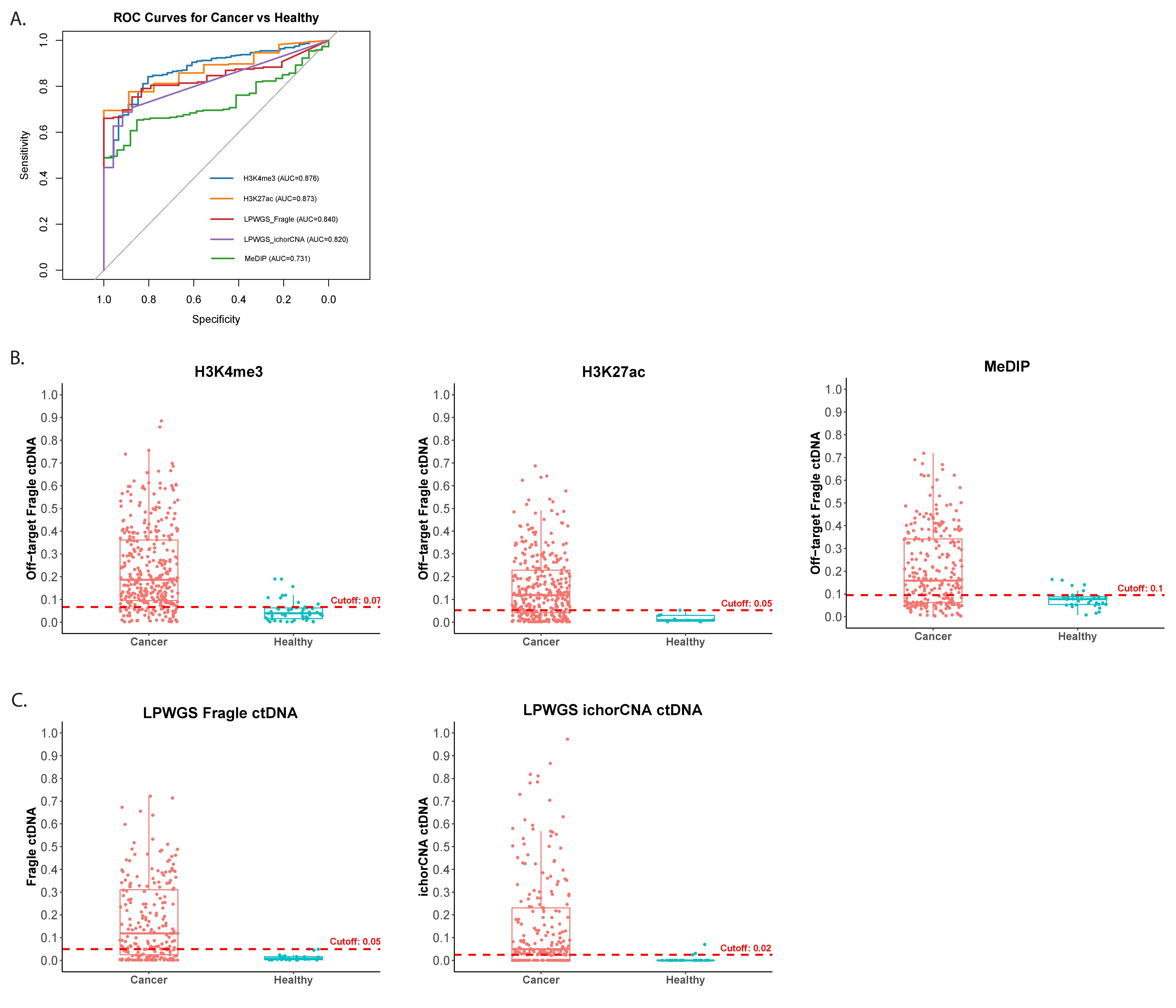
**Supplementary Figure 9.** **A.** ROC curves for using ctDNA to distinguish cancer plasma from healthy plasma samples; **B.** Fragle ctDNA comparison between cancer plasma and healthy plasma samples across H3K4me3 cfChIP-seq, H3K27ac cfChIP-seq, and cfMeDIP assays; **C.** LPWGS Fragle ctDNA and ichorCNA ctDNA comparisons between cancer plasma and healthy plasma samples.

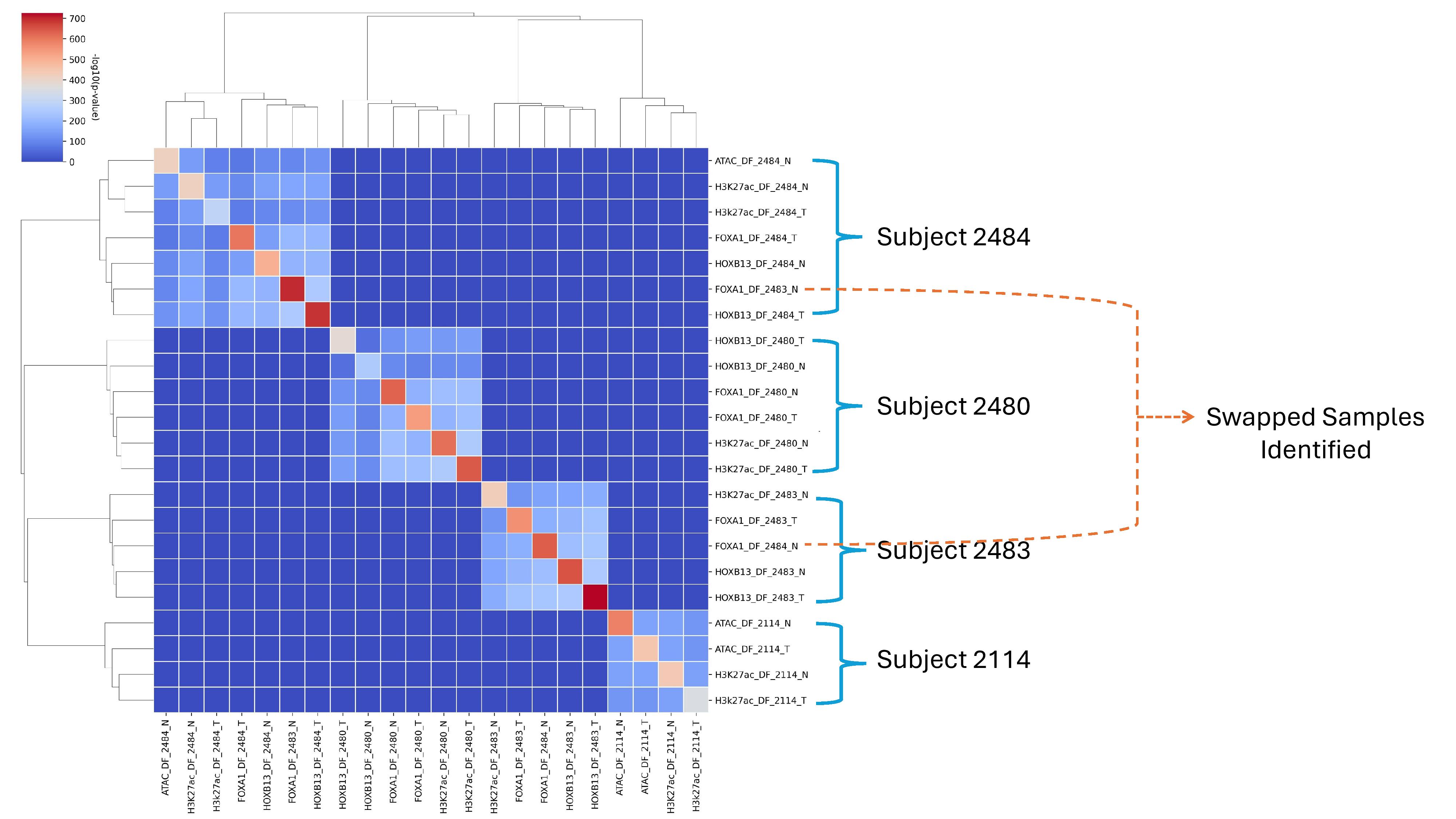

**Supplementary Figure 10.** *SMaSH* SNAP fingerprint clustering identifies mismatched ChIP-seq samples from GSE130408^2^.

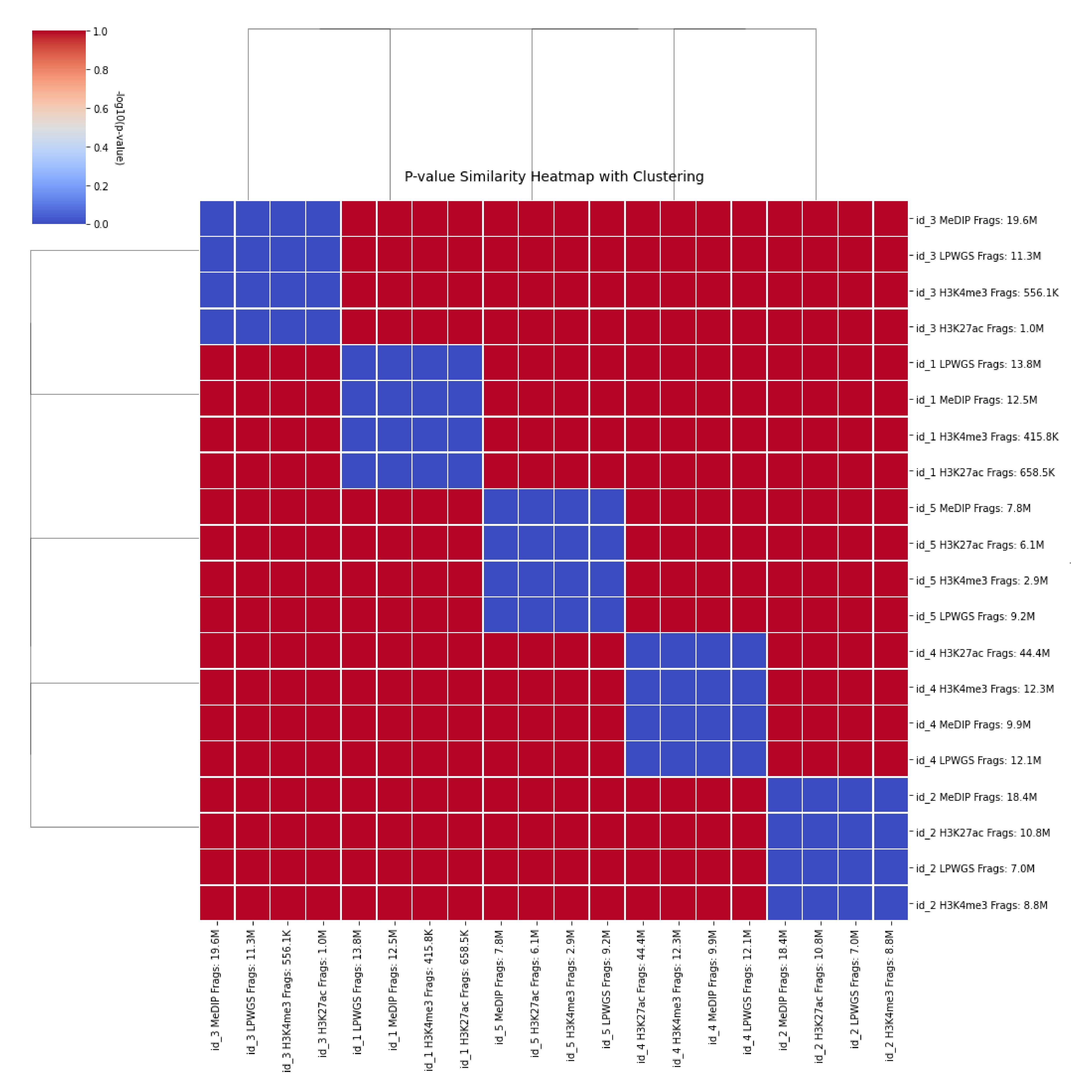

**Supplementary Figure 11.** *SMaSH* SNP fingerprinting for MeDIP, LPWGS, H3K4me3, and H3K27ac samples from the same individuals.

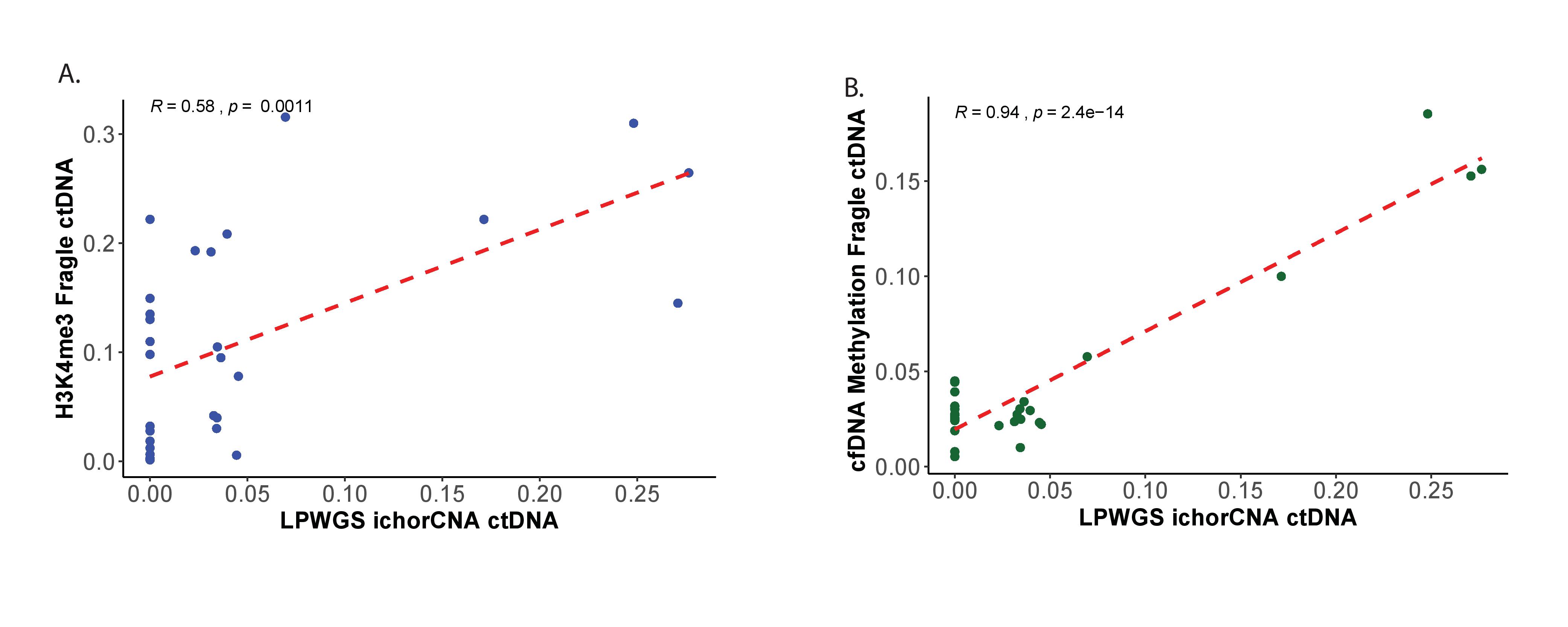

**Supplementary Figure 12.** Osteosarcoma plasma samples: **A.** Correlation between LPWGS ichorCNA-estimated ctDNA and H3K4me3 Fragle-estimated ctDNA; **B.** Correlation between LPWGS ichorCNA-estimated ctDNA and cfDNA methylation Fragle-estimated ctDNA.

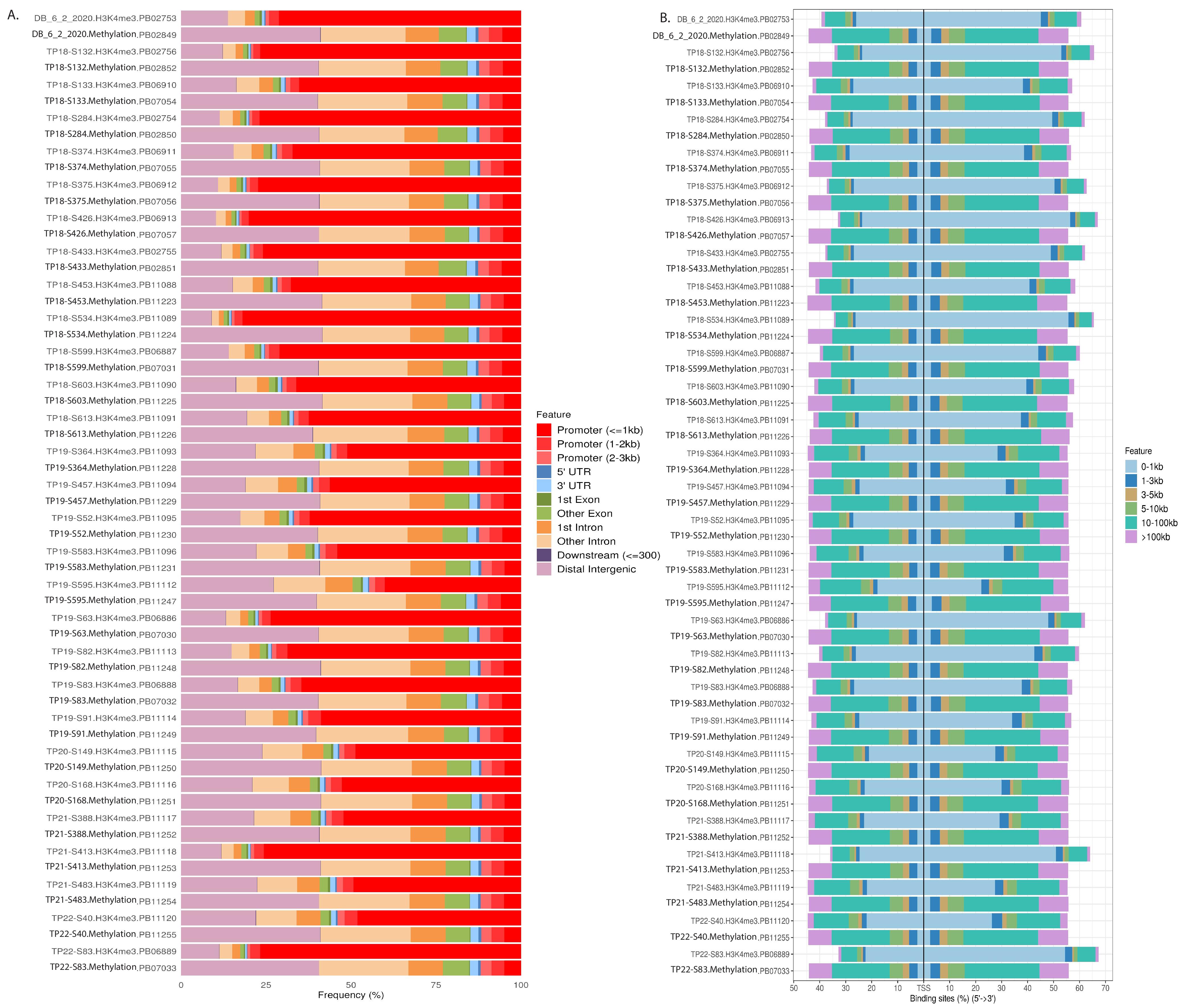

**Supplementary Figure 13.** Osteosarcoma plasma H3K4me3 cfChIP-seq and cfDNA methylation peak signal distribution in **A.** genomic locations and **B.** transcription start site locations.

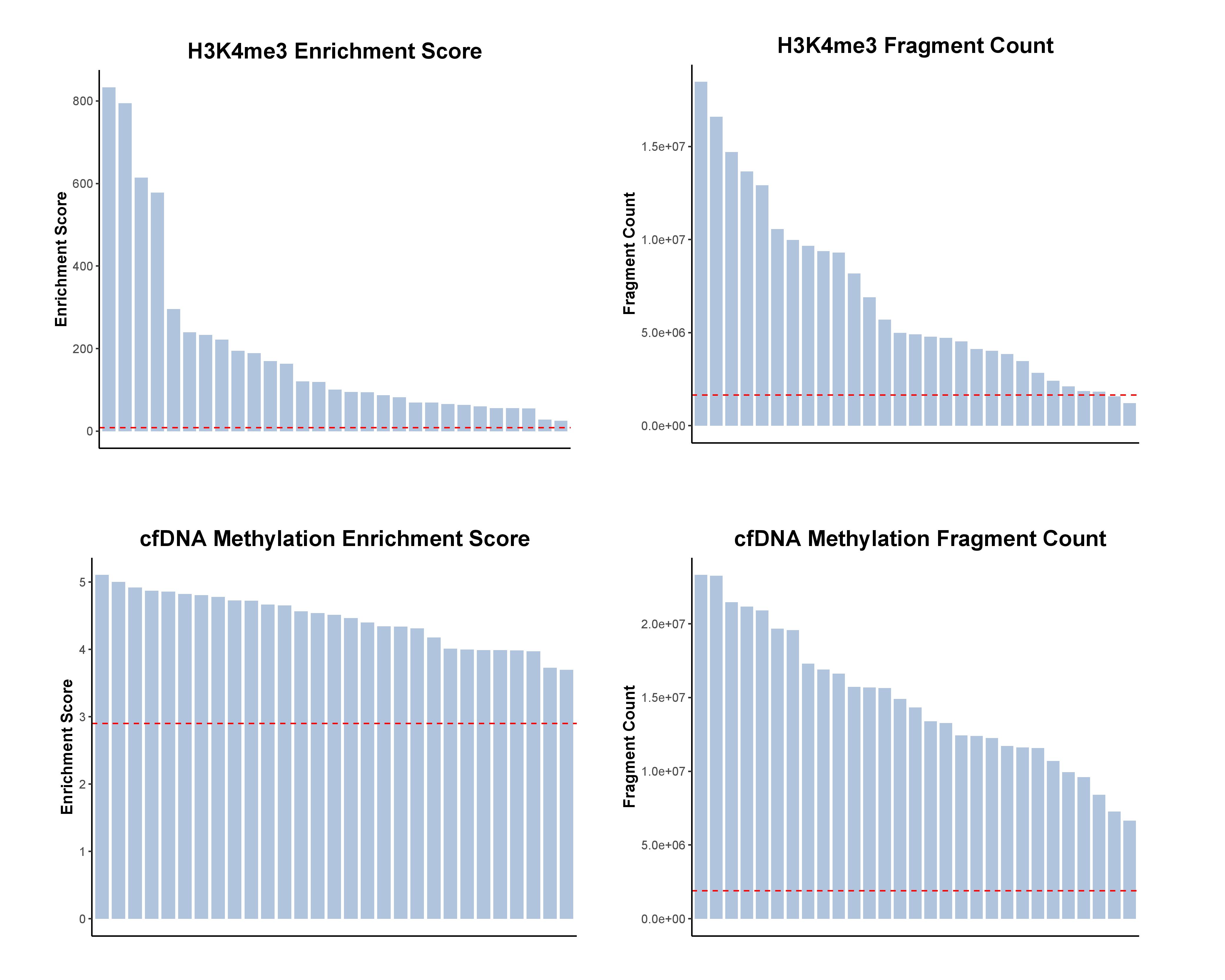

**Supplementary Figure 14.** Quality control metrics for osteosarcoma plasma H3K4me3 cfChIP-seq and cfDNA methylation samples. Red dashed lines represent established quality control cut-offs (cell-free chromatin H3K4me3: enrichment score > 8.7 and fragment count > 1.64e+06; cfDNA methylation: enrichment score > 2.9 and fragment count > 1.2e+06).

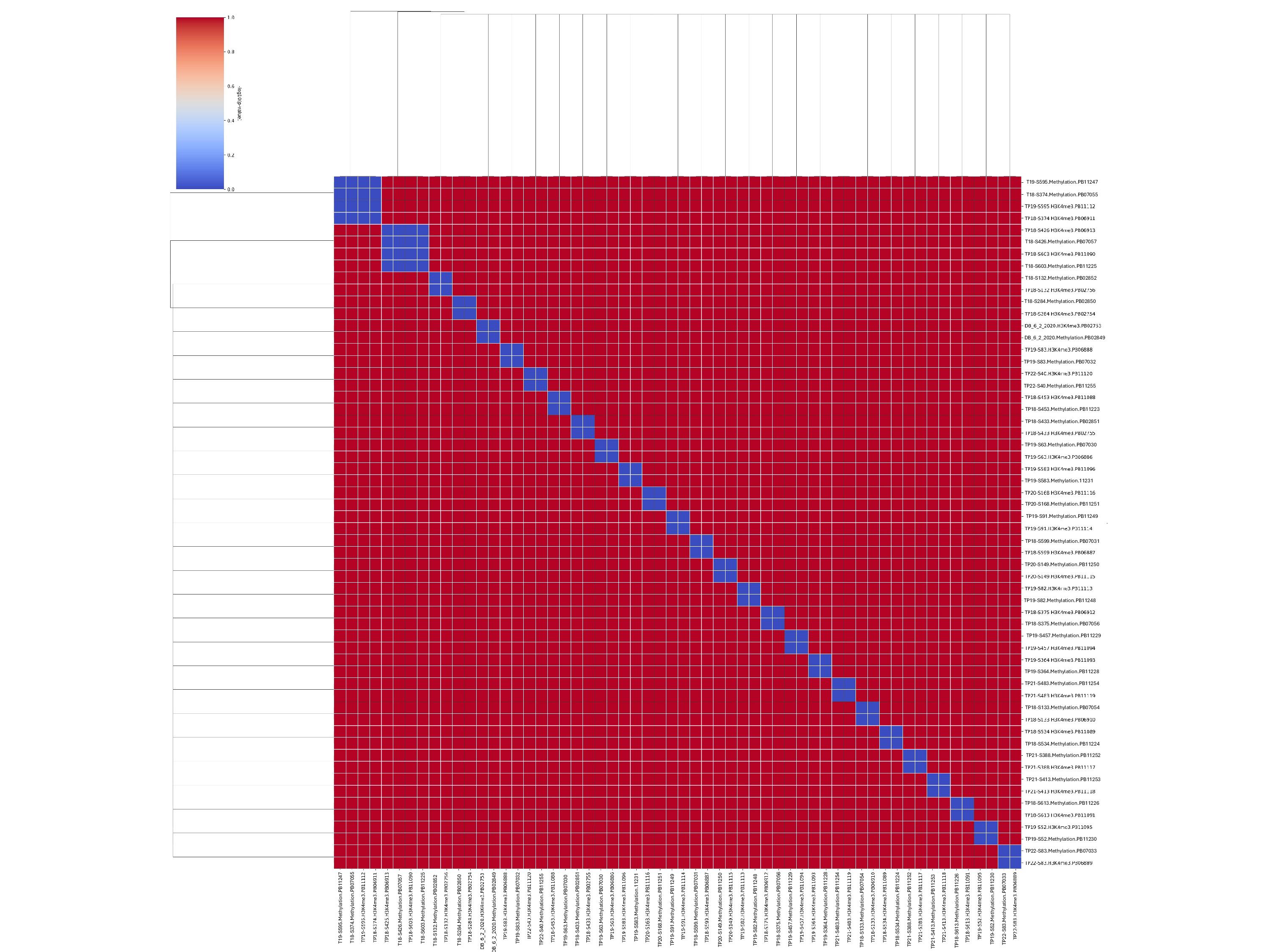

**Supplementary Figure 15.** *SMaSH* SNP fingerprinting for osteosarcoma plasma H3K4me3 cfChIP-seq and cfDNA methylation samples.

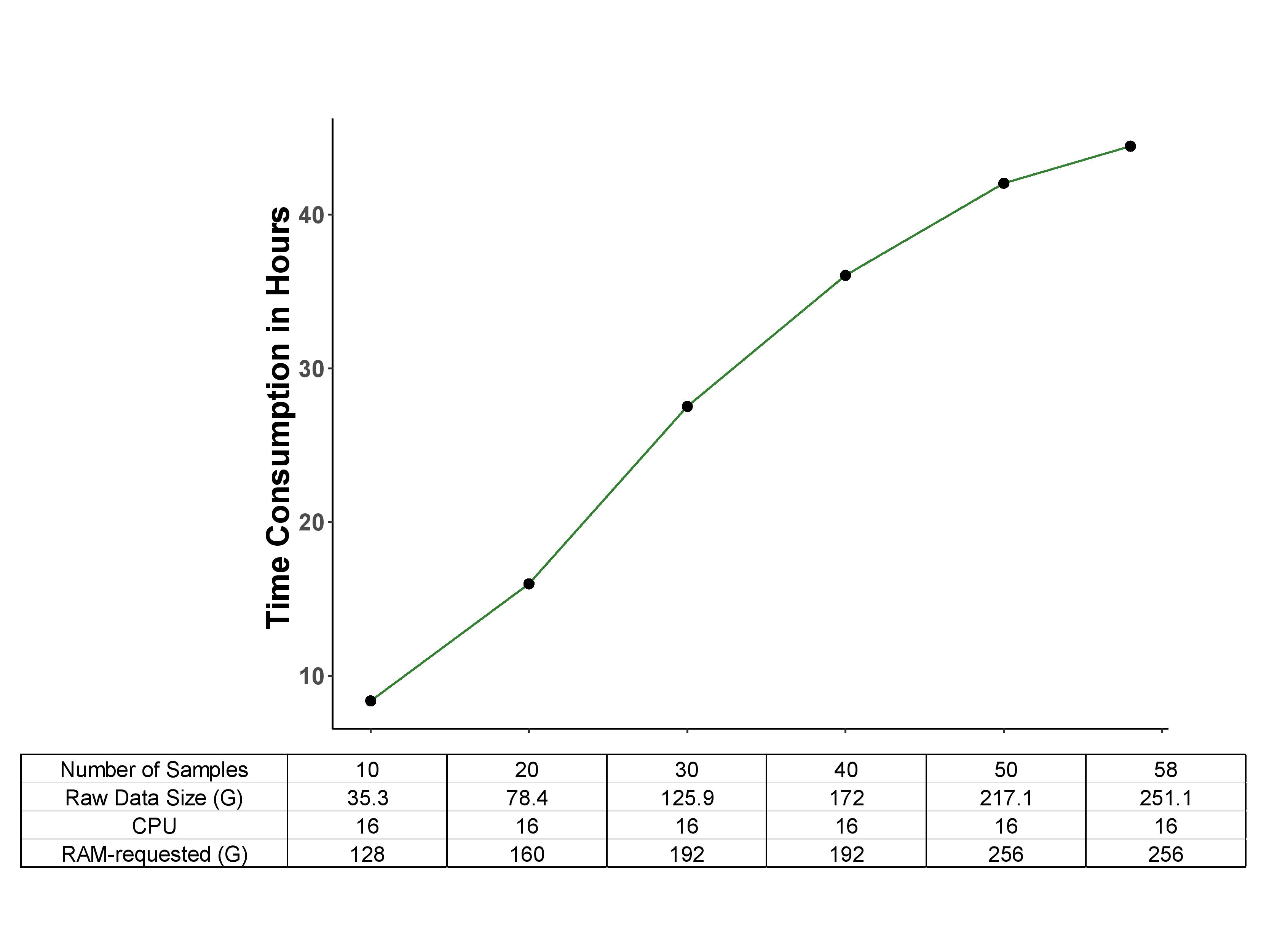
**Supplementary Figure 16.** Computational performance and resource usage of SNAP on the osteosarcoma cohort. All analyses were performed on a workstation equipped with an AMD Ryzen Threadripper PRO 5975WX processor (32 physical cores, 64 logical cores) and 503 GB RAM, running Ubuntu 24.04.3 LTS.
